# An integrated single-cell and spatial transcriptomic atlas of thyroid cancer progression identifies prognostic fibroblast subpopulations

**DOI:** 10.1101/2025.01.08.631962

**Authors:** Matthew A. Loberg, George J. Xu, Sheau-Chiann Chen, Hua-Chang Chen, Claudia C. Wahoski, Kailey P. Caroland, Megan L. Tigue, Heather A. Hartmann, Jean-Nicolas Gallant, Courtney J. Phifer, Andres Ocampo, Dayle K. Wang, Reilly G. Fankhauser, Kirti A. Karunakaran, Chia-Chin Wu, Maxime Tarabichi, Sophia M. Shaddy, James L. Netterville, Sarah L. Rohde, Carmen C. Solorzano, Lindsay A. Bischoff, Naira Baregamian, Barbara A. Murphy, Jennifer H. Choe, Jennifer R. Wang, Eric C. Huang, Quanhu Sheng, Luciane T. Kagohara, Elizabeth M. Jaffee, Ryan H. Belcher, Ken S. Lau, Fei Ye, Ethan Lee, Vivian L. Weiss

## Abstract

Thyroid cancer progression from curable well-differentiated thyroid carcinoma to highly lethal anaplastic thyroid carcinoma is distinguished by tumor cell de-differentiation and recruitment of a robust stromal infiltrate. Combining an integrated thyroid cancer single-cell sequencing atlas with spatial transcriptomics and bulk RNA-sequencing, we define stromal cell subpopulations and tumor-stromal cross-talk occurring across the histologic and mutational spectrum of thyroid cancer. We identify distinct inflammatory and myofibroblastic cancer-associated fibroblast (iCAF and myCAF) populations and perivascular-like populations. The myCAF population is only found in malignant samples and is associated with tumor cell invasion, *BRAF*^V600E^ mutation, lymph node metastasis, and disease progression. Tumor-adjacent myCAFs abut invasive tumor cells with a partial epithelial-to-mesenchymal phenotype. Tumor-distant iCAFs infiltrate inflammatory autoimmune thyroid lesions and anaplastic tumors. In summary, our study provides an integrated atlas of thyroid cancer fibroblast subtypes and spatial characterization at sites of tumor invasion and de-differentiation, defining the stromal reorganization central to disease progression.

## Main

In 1986, Harold Dvorak identified tumors as “wounds that do not heal” in recognition of the inflamed tumor microenvironment undergoing persistent extracellular matrix remodeling that resembles chronic wound healing.^1^ Fibroblasts, a central stromal component of wound healing, are highly plastic cells that become activated to remodel the extracellular matrix in response to tissue injury.^2^ Thus, in the chronic wound state of tumors, cancer-associated fibroblasts (CAFs) play a critical role in modulating the tumor microenvironment and tumor cell phenotypes.

Studies across solid tumors have implicated CAFs as supporting tumor progression and repressing anti- tumor immunity, making them promising therapeutic targets.^3^ However, attempts to broadly deplete CAFs in mouse models have resulted in aggressive tumor growth, highlighting the complex and tumor- restrictive properties of CAFs.^4,5^ Clinical trials targeting CAFs or extracellular matrix have also had limited success.^6^ To improve CAF targeting and modulation strategies, multiple groups have sought to identify distinct tumor-promoting CAF subpopulations and functions. Single-cell sequencing efforts across tumor types have now defined numerous CAF subtypes.^7–15^ Broad CAF subtypes, including myofibroblast (myCAF), inflammatory (iCAF), and antigen-presenting are commonly conserved across tumor types. However, the plastic and heterogenous CAF functionality recruited during tumor progression remains poorly defined, with subsets such as myCAFs serving critical roles in both matrix remodeling and immune modulation.^12,16^ Additionally, several studies have identified perivascular populations as potential CAF precursors with CAF-like gene expression profiles.^10,11,14^ Despite identification of numerous CAF subtypes, CAFs remain an elusive clinical target. A better understanding of CAF phenotypes and plasticity within disease progression is needed to improve cancer therapeutics.

Well-differentiated thyroid cancers (WDTCs) are associated with excellent prognosis, yet in rare instances clinically progress to anaplastic thyroid carcinoma (ATC),^17,18^ a carcinoma with an abysmal median survival of 3-5 months.^19^ Despite the rarity of WDTC clinical progression, ATCs often contain co- occurring, adjacent WDTC histologic regions, suggesting pathologic progression of WDTC to ATC.^20–24^ This spatial progression is accompanied by a robust accumulation of tumor stroma.^25^ The stark difference in tumor aggression and stromal infiltrate between WDTC and ATC, as well as their co-occurrence in ATC, make thyroid cancer an excellent model for tracking CAF phenotypes over spatial progression from curable to lethal disease. In our previous work, we developed a gene expression signature, the molecular aggression and prediction (MAP) score, that implicated fibroblasts as enriched in thyroid cancers at risk for disease recurrence, progression, or de-differentiation to ATC.^25^ CAFs have previously been identified in thyroid cancer single-cell sequencing studies,^26–29^ yet a comprehensive profiling of CAF phenotypes over spatial progression is lacking.

In this study, we integrate single-cell RNA-sequencing data from 81 published thyroid cancer samples,^26–32^ generating a single-cell atlas to identify CAF subtypes across the landscape of tumor histology.

Additionally, we perform spatial transcriptomics on 28 thyroid tumors to investigate CAF locations and interactions during thyroid cancer progression. Remarkably, we include seven samples with co-occurring papillary (WDTC) and anaplastic histology, providing a unique evaluation of the spatial progression of thyroid cancer. Finally, we map CAF subtypes to four large thyroid cancer bulk RNA-sequencing cohorts, demonstrating their potential prognostic role in metastasis and disease progression.^31,33^ These data identify myCAFs as a potential biomarker for aggressive thyroid tumors and highlight their phenotype and tumor cell-interactions during disease progression.

## Results

### Generation of thyroid cancer single-cell atlas

We integrated published thyroid cancer single-cell RNA-sequencing samples from seven papers with samples across the histologic and mutational landscape of thyroid cancer progression, including 20 normal thyroids/paratumors, 39 papillary thyroid cancers (PTC, the most common WDTC), and 22 ATCs (Fig. 1a and Supplementary Table 1). ^26–32^ In total, 423,733 cells were integrated and clustered to identify broad cell populations (Fig. 1b). Tumor, normal, and microenvironment populations expressing canonical marker genes were identified across samples from all seven papers, including stromal cells with expression of fibroblast markers *DCN* and *COL1A1* (Fig. 1c-e and Extended Data Fig. 1a-c). Most paratumors (15/17, 88%) had a low frequency (0-7%) of PTC cells, consistent with their tumor-adjacent location. We further confirmed that thyrocyte and tumor cell populations had expected gene expression patterns by mapping previously developed thyroid differentiation (TDS) and tumors scores.^29,33^ TDS showed the highest enrichment in thyrocytes, a gradient in PTC cells, and no enrichment in ATC, consistent with de-differentiation as tumors progress. *BRAF* score, designed to identify *BRAF*^V600E^ PTCs,^33^ was highest in the PTC cluster. A 12 gene signature designed to distinguish ATC tumor cells from PTC tumor cells^29^ identified both ATC cells as well as stromal cells, in line with the de-differentiated mesenchymal phenotype of ATC tumor cells (Extended Data Fig. 1d,e). Altogether, we generated an integrated thyroid cancer atlas that includes samples across a spectrum of thyroid cancer de- differentiation.

**Fig. 1:**
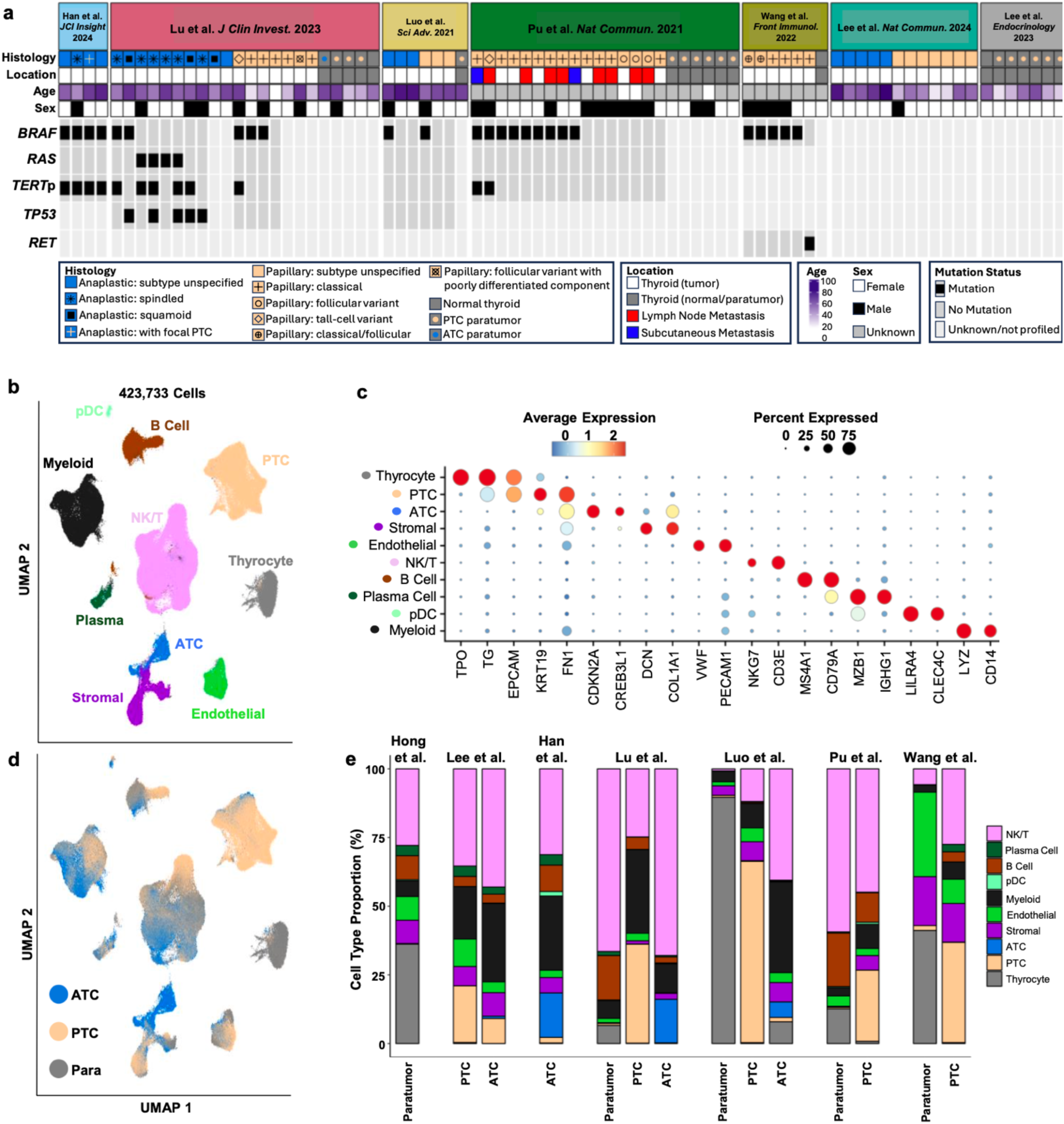
Integrated single-cell atlas of thyroid cancer progression. **a**, Oncoplot showing histology, location, age, and sex meta data (top) and key driver mutations (bottom) for thyroid cancer publicly available single-cell RNA-sequencing samples. **b**, Uniform Manifold Approximation and Projection (UMAP) plot depicting the 423,733 cells in the integrated thyroid cancer single-cell atlas labeled by broad cell type. **c**, Scaled dot plot showing canonical markers for broad single-cell populations from **b**. **d**, UMAP plot of the integrated single-cell atlas colored by tumor histology with broad groupings of anaplastic thyroid carcinoma (ATC, blue), papillary thyroid carcinoma (PTC, light orange), or paratumor/normal (Para, grey). **e**, Bar plots showing overall broad cell type composition for each paper in the single-cell atlas split by tumor histology group (Paratumor, PTC, ATC). Abbreviations: pDC, plasmacytoid dendritic cell; NK/T, natural killer/T cell.

### Identification of CAF and perivascular subpopulations

To identify fibroblast subpopulations, we performed subclustering of 24,463 stromal cells in the single-cell atlas and identified six main clusters (Fig. 2a). We proceeded with analysis of clusters 0-4, as cluster 5 was largely from one metastatic lymph node sample from Pu et al. (Fig 2b,c and Extended Data Fig. 2a). We confirmed broad expression of *VIM* and *PDGFRB* across clusters, consistent with their mesenchymal phenotype (Fig. 2d). We observed expression of perivascular markers *RGS5* and *NOTCH3* in clusters 4, 3, and 0 whereas pan-fibroblast markers *DCN* and *PDGFRA* were expressed mainly by clusters 1 and 2, with some expression in cluster 4 (Fig. 2e,f). We thus hypothesized that our stromal subclustering identifies both CAF and perivascular populations. To better classify these populations, we generated Seurat module scores^34^ for published CAF and perivascular marker genes from diverse tumor types.^8–15,35,36^ Amongst hypothesized CAFs, cluster 0 showed enrichment of each iCAF gene set tested, whereas cluster 2 showed higher scores for matrix CAFs, myCAFs, and TGFβ- CAF gene sets. Within the hypothesized perivascular cell clusters, there was broad enrichment of perivascular gene sets, with cluster 0 having higher module scores for smooth muscle cells (SMCs) and cluster 3 having higher scores for pericyte gene sets. Cluster 4 showed enrichment for all perivascular gene sets but had no previously defined phenotype from the literature (Fig. 2g,h). We next performed differential gene expression analysis to identify marker genes of each stromal subcluster (Supplementary Table 2 and Extended Data Fig. 2b). Top marker genes of clusters included known iCAF marker genes *CXCL12* and *APOD* (cluster 0), the myCAF gene *POSTN* and the activated CAF gene *FAP* (cluster 2), *APOE* (cluster 4), pericyte markers *HIGD1B* and *COX4I2* (cluster 0), and SMC markers *ACTA2* and *TAGLN* (cluster 3). In addition to marking SMCs, *ACTA2* and *TAGLN* have been described as myCAF markers^13^ and were also expressed by cluster 2 (Fig. 2i-k). Based on the concordance of published gene sets and our differential gene expression analysis, we labeled cluster 1 as iCAFs, cluster 2 as myCAFs, and cluster 0 as pericytes. Because cluster 3 expressed perivascular and SMC markers, we labeled it as vascular SMCs (vSMCs). We labeled cluster 3 as APOE+ perivascular-like (APOE+ PVL) based on its moderate expression of perivascular marker genes (Fig. 2l). Gene ontology analysis of the marker genes for each population highlighted the broad roles of each population in matrix remodeling (Extended Data Fig. 2c). Analysis of signaling pathway activation highlighted known fibroblast activating pathways TGFβ and Wnt signaling in myCAFs, estrogen and androgen signaling in iCAFs, and tumor necrosis factor- related apoptosis-induced ligand (TRAIL) signaling in perivascular populations, consistent with the recently established role of TRAIL in facilitating cross-talk between endothelial cells and perivascular cells (Extended Data Fig. 2d).^37^ In conclusion, we identified myCAF, iCAF, and perivascular subpopulations that align with published gene sets for these populations.

**Fig. 2:**
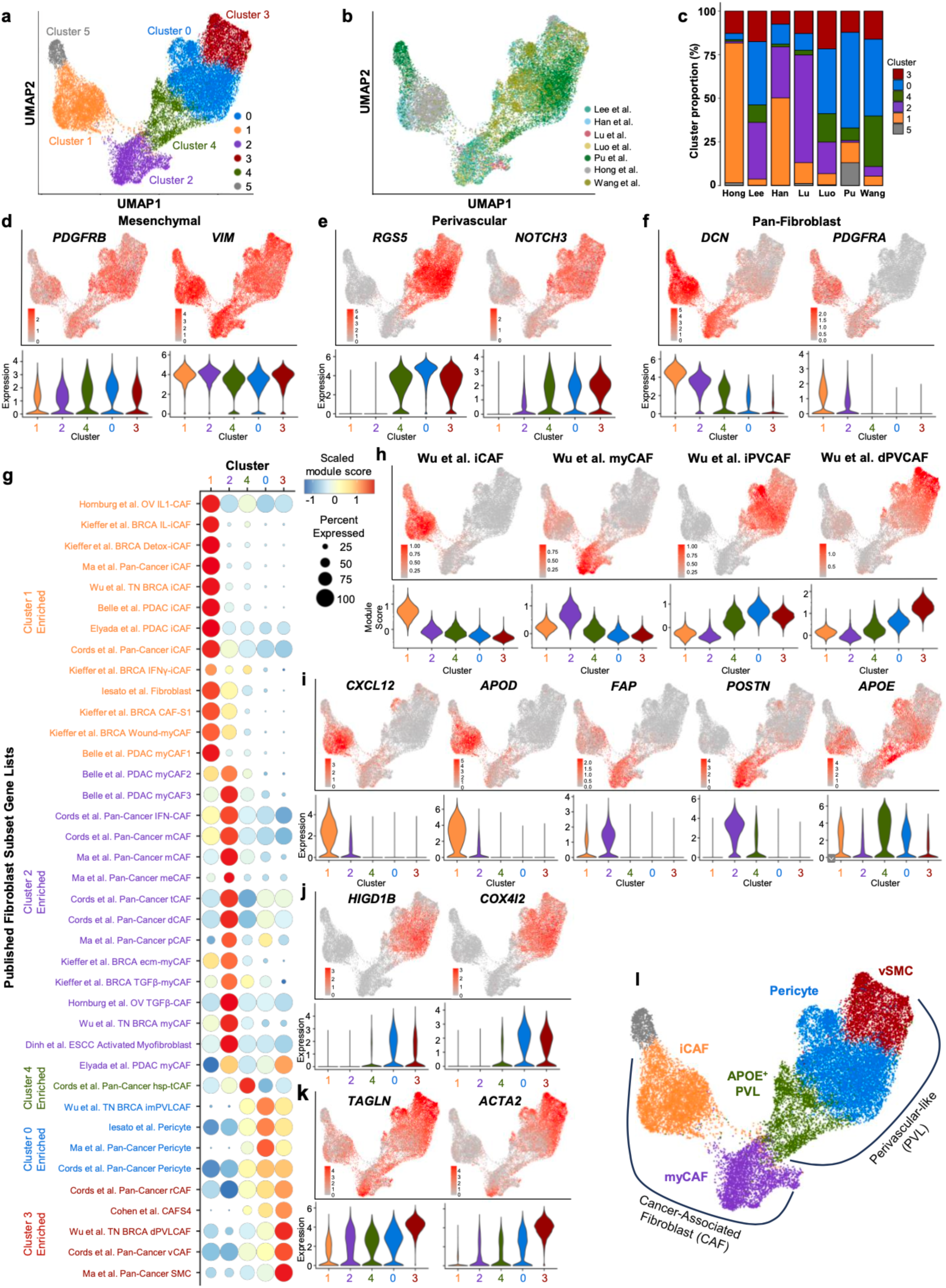
Stromal subclustering identifies key CAF and perivascular populations. a,b, Uniform Manifold Approximation and Projection (UMAP) showing subclustering of the broad stromal cluster from Fig. 1b colored by (**a**) cluster and (**b**) paper. **c**, Bar plots showing cluster proportion for each paper. **d-f**, UMAP of stromal subclustering depicting expression of (**d**) mesenchymal marker genes *PDGFRB* and *VIM*, (**e**) perivascular marker genes *RGS5* and *NOTCH3*, and (**f**) pan-fibroblast markers *DCN* and *PDGFRA* with violin plots below each UMAP showing expression by cluster. **g**, Scaled dot plot showing module scores for published cancer-associated fibroblast (CAF) and perivascular populations by stromal subcluster. **h**, UMAP of stromal subclustering showing module scores for CAF (inflammatory CAF, iCAF; myofibroblast CAF, myCAF) and perivascular-like (immature perivascular-like CAF, iPVCAF; differentiated perivascular-like CAF, dPVCAF) populations from Wu et al. with violin plots below depicting expression by stromal subcluster. **i-k**, UMAP showing marker genes of stromal subclusters with violin plots below. **l**, Diagram outlining labeling of stromal subclustering into broad CAF and perivascular-like (PVL) groups with identification of individual clusters as iCAF, myCAF, APOE+ PVL, pericyte, and vascular smooth muscle cell (vSMC). Abbreviations: OV, ovarian cancer; BRCA, breast cancer; TN BRCA, triple negative breast cancer; PDAC, pancreatic ductal adenocarcinoma; ESCC, esophageal squamous cell carcinoma; mCAF, matrix CAF; meCAF, metabolic CAF; tCAF, tumor-like CAF; dCAF, dividing CAF; ecmCAF, extracellular matrix CAF; hsp-tCAF, heat-shock protein tumor-like CAF.

### myCAFs are associated with aggressive tumors

We next sought to determine the association of stromal subclusters with tumor histology and disease progression. We observed that myCAFs were predominantly found in malignant samples, with a subset of ATCs having the highest proportion of myCAFs (Fig. 3a,b; Extended Data Fig. 3a). Differential abundance testing confirmed enrichment of myCAFs in malignant samples and ATCs and identified a robust pericyte signal in PTC. iCAFs showed enrichment in paratumors and in ATC relative to PTC (Fig 3c). iCAFs are commonly thought to be tumor-distant,^38^ aligning with their presence in paratumor sampling. As previous studies have associated CAFs and desmoplastic stroma with *BRAF*^V600E^ PTCs,^25,39^ we scored PTCs based on expression of genes from The Cancer Genome Atlas (TCGA) *BRAF*-*RAS* score (BRS).^33^ PTCs with myCAFs had higher median *BRAF* scores than PTCs without myCAFs (p=0.017) and PTCs with high median *BRAF* scores showed broad enrichment of stromal populations, including myCAFs (Fig. 3d,e). To determine how myCAF phenotypes shift as tumors de-differentiate from PTC to ATC, we performed differential gene expression analysis of myCAFs based on sample histology. myCAFs in PTC showed a classic myofibroblast phenotype identified in pancreatic cancer,^13,38^ with *ACTA2*, *TAGLN,* and collagen gene expression. *RGS5* was also upregulated in PTC myCAFs, suggesting a potential pericyte precursor for this population, consistent with the abundance of pericytes observed in differential abundance analysis. In contrast, ATC myCAFs exhibited upregulation of immune modulating genes (Fig. 3f).

**Fig. 3:**
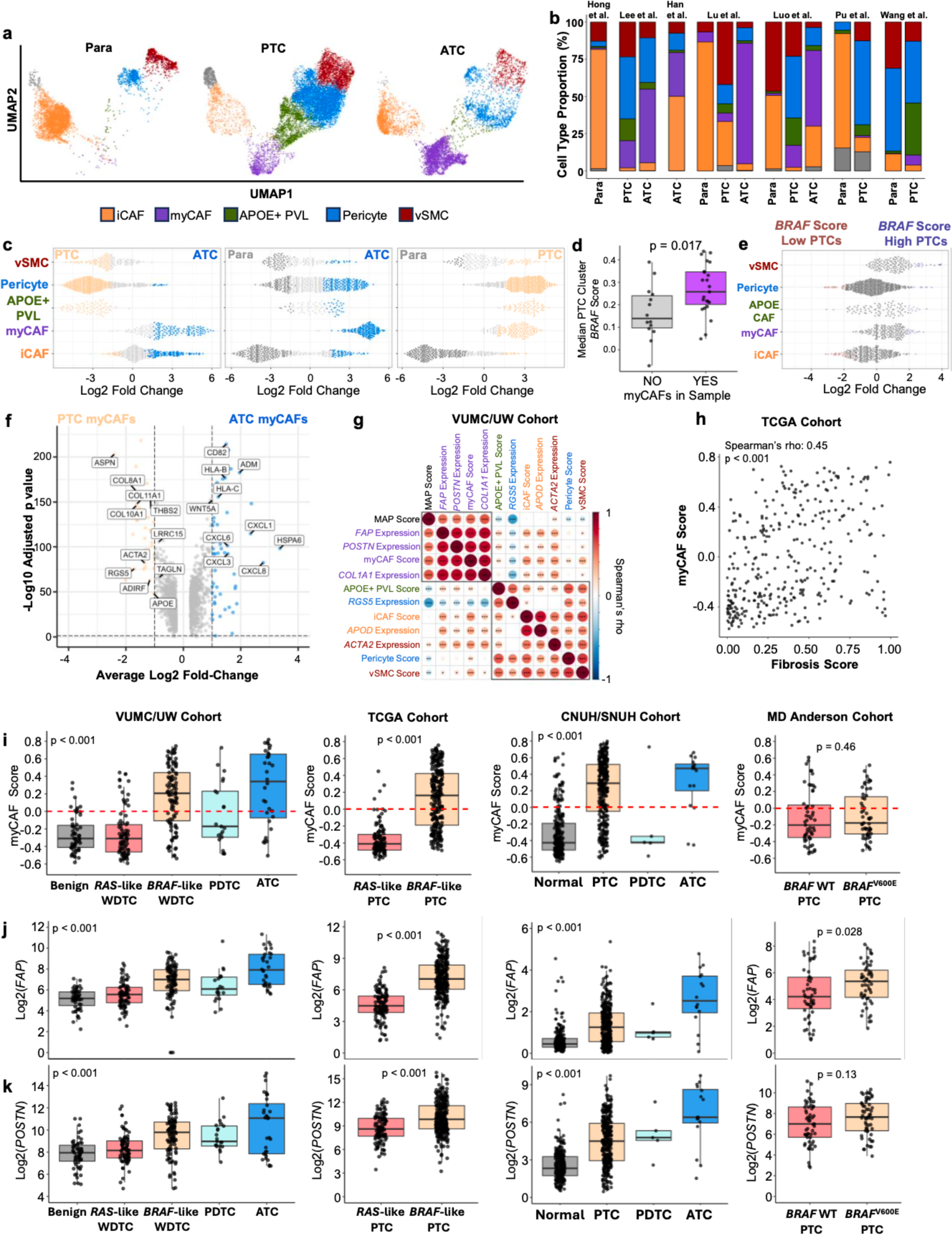
myCAFs are upregulated in malignant tumors across thyroid cancer sequencing cohorts. **a**, Stromal subclustering Uniform Manifold Approximation and Projection (UMAP) split by tumor histology with broad groupings of paratumor/normal (Para, left), papillary thyroid cancer (PTC, middle), and anaplastic thyroid carcinoma (ATC, right). **b**, Bar plots showing cell type proportion for each tumor histology grouping within each paper. **c**, Milo differential abundance testing of cancer-associated fibroblast (CAF) and perivascular stromal cell populations (inflammatory CAF, iCAF; myofibroblast CAF, myCAF; APOE+ perivascular-like, APOE+ PVL; pericyte; vascular smooth muscle cell, vSMC) between PTC and ATC samples (left), paratumor/normal and ATC samples (middle), and paratumor/normal and PTC samples (right). Individual dots depict neighborhoods calculated by Milo. Coloring of individual neighborhoods as dark grey (normal/Paratumor), light orange (PTCs), or blue (ATC) indicates differential abundance with a spatial false discovery rate (FDR) of less than 0.1. Neighborhoods colored light grey have a spatial FDR greater than 0.1. **d**, Boxplot depicting the median *BRAF* module score of PTC cells for individual PTC samples split by whether the sample contains myCAFs. p-value calculated with Wilcoxon rank sum test. **e**, Milo differential abundance testing of CAF and perivascular stromal cell populations (iCAF, myCAF, APOE+ PVL, pericyte, vSMC) between PTC tumors split based on *BRAF* score high (median *BRAF* score for PTC cells within the tumor greater than 50^th^ percentile) or *BRAF* score low (median *BRAF* score for PTC cells within the tumor less than 50^th^ percentile). Coloring of individual neighborhoods as red (BRAF score low) or blue (BRAF score high) indicates differential abundance with a spatial FDR of less than 0.1. Neighborhoods colored grey have a spatial FDR greater than 0.1. **f,** Volcano plot showing differentially expressed genes between PTC myCAFs (left, light orange) and ATC myCAFs (right, blue). Colored dots indicate genes with an absolute log2 fold-change of at least 1.0 and adjusted p<0.05. **g,** Corrplot showing Spearman’s rho correlations between single-sample GSVA (ssGSVA) scores for stromal subpopulations, ssGSVA score for molecular aggression and prediction (MAP) score, and expression of marker genes for stromal subpopulations in the Vanderbilt University/University of Washington bulk RNA-sequencing cohort (VUMC/UW cohort). Axes are ordered by hierarchical clustering. Boxes indicate hierarchical clustering groups. Significance levels indicate *p<0.05, **p<0.01, or ***p<0.001. **h,** Spearman’s rho correlation of myCAF ssGSVA score and a fibrosis score derived from hematoxylin and eosin imaging data of The Cancer Genome Atlas PTCs (TCGA cohort).^39^ **i,** Boxplots showing myCAF score by diagnosis across four distinct bulk RNA-sequencing cohorts. **j,k**, Boxplots showing log2 transformed expression of CAF marker genes (**j**) *FAP* and (**k**) *POSTN* by diagnosis across four distinct bulk RNA-sequencing cohorts. P-values for **i-k** calculated with Wilcoxon rank-sum test. Kruskal-Wallis test with subsequent pairwise Wilcoxon rank-sum tests with Bonferroni’s correction was used when comparing more than two groups. Abbreviations: WDTC, well-differentiated thyroid cancer; PDTC, poorly differentiated thyroid cancer, CNUH/SNUH, Chungnam/Seoul National University Hospitals.

To validate the association of myCAFs with aggressive tumors, we used single-sample gene set variation analysis (GSVA) to predict abundance of CAF and perivascular subpopulations in four large thyroid cancer bulk RNA-sequencing cohorts.^25,31,33^ We correlated stromal GSVA scores with our previously published MAP score, which identifies aggressive, fibroblast-rich tumors.^25^ Across cohorts, MAP score was correlated with myCAF GSVA scores and the expression of myCAF-associated genes (Fig. 3g, Extended Data Fig. 3b). To validate that myCAFs are enriched in tumors with increased stroma and matrix remodeling, we identified a positive correlation (Spearman’s rho 0.45, p<0.001) between myCAF score in the TCGA cohort and a fibrosis score previously generated from histology imaging data (Fig. 3h).^39^ Next, we assessed the predicted abundance of stromal cells across benign, WDTC, and ATC in these cohorts. myCAF scores and myCAF marker genes were upregulated in *BRAF*-like WDTCs and ATCs across Vanderbilt University/University of Washington (VUMC/UW), TCGA, and Chungnam/Seoul National University Hospitals (CNUH/SNUH) cohorts. Aggressive PTCs from MD Anderson showed increased *FAP* expression in *BRAF*^V600E^ tumors (p=0.028) (Fig. 3i-k). In contrast, iCAFs and perivascular subpopulations did not show clear shifts in predicted abundance with tumor de-differentiation (Extended Data Fig. 3c-f). iCAFs were increased in VUMC/UW ATCs relative to WDTCs (p=0.0019) and in *BRAF*- like WDTCs in the TCGA cohort relative to *RAS*-like WDTCs (p<0.001), but the CNUH/SNUH cohort showed the highest predicted iCAF levels in benign thyroids. However, many benign and malignant surgically removed thyroids have background autoimmune inflammatory processes (e.g., Hashimoto thyroiditis). To assess whether increased iCAF infiltrate in benign thyroids was related to autoimmune disease, we evaluated the predicted abundance of iCAFs in benign lesions from the VUMC/UW cohort that underwent detailed pathologic and clinical evaluation. Amongst VUMC/UW benign thyroids, there was a higher predicted abundance of iCAFs (p=0.0015), but not myCAFs, in patients with confirmed Hashimoto thyroiditis, as compared to non-autoimmune benign follicular nodular disease (Extended Data Fig. 3g). These data indicate that autoimmune conditions may have fibroblasts with inflammatory phenotypes similar to iCAFs.

We next assessed the association of stromal populations with features of aggressive disease and patient outcomes. In malignant samples in the VUMC/UW cohort, myCAF score was associated with worse progression-free (PFS) and overall survival (OS; Fig. 4a,b). We also observed worse PFS in myCAF score high tumors in a cohort of aggressive PTCs (p=0.045, Fig. 4c), indicating that decreases in survival in myCAF-rich tumors are not solely driven by ATC. The TCGA cohort was designed to consist of indolent PTCs with few survival events. However, we identified that TCGA “intermediate- or high-risk” PTCs, as designated by the 2009 ATA guidelines,^40^ had higher myCAF scores than “low-risk” PTCs (p<0.001, Fig. 4d). Additionally, in both TCGA and VUMC/UW cohorts, tumors with extrathyroidal extension (p<0.001 TCGA; p<0.001 VUMC/UW) or lymph node metastases (p<0.001 TCGA; p<0.001 VUMC/UW) had higher myCAF scores (Fig. 4e,f). Increased myCAF score in primary tumors with distant metastases was also observed in the VUMC/UW cohort (p=0.034) and lymph node metastases had higher myCAF scores than primary malignant tumors in the VUMC/UW cohort (p=0.036, Fig. 4g,h). Finally, PTCs with aggressive histologic subtypes tall cell or diffuse sclerosing had the highest myCAF scores (p<0.001, Fig. 4i). In total, we identified myCAFs as enriched in malignant samples and ATCs across our single-cell atlas and identified association with ATC, PFS, OS, and aggressive WDTC features in bulk RNA-sequencing cohorts.

**Fig. 4:**
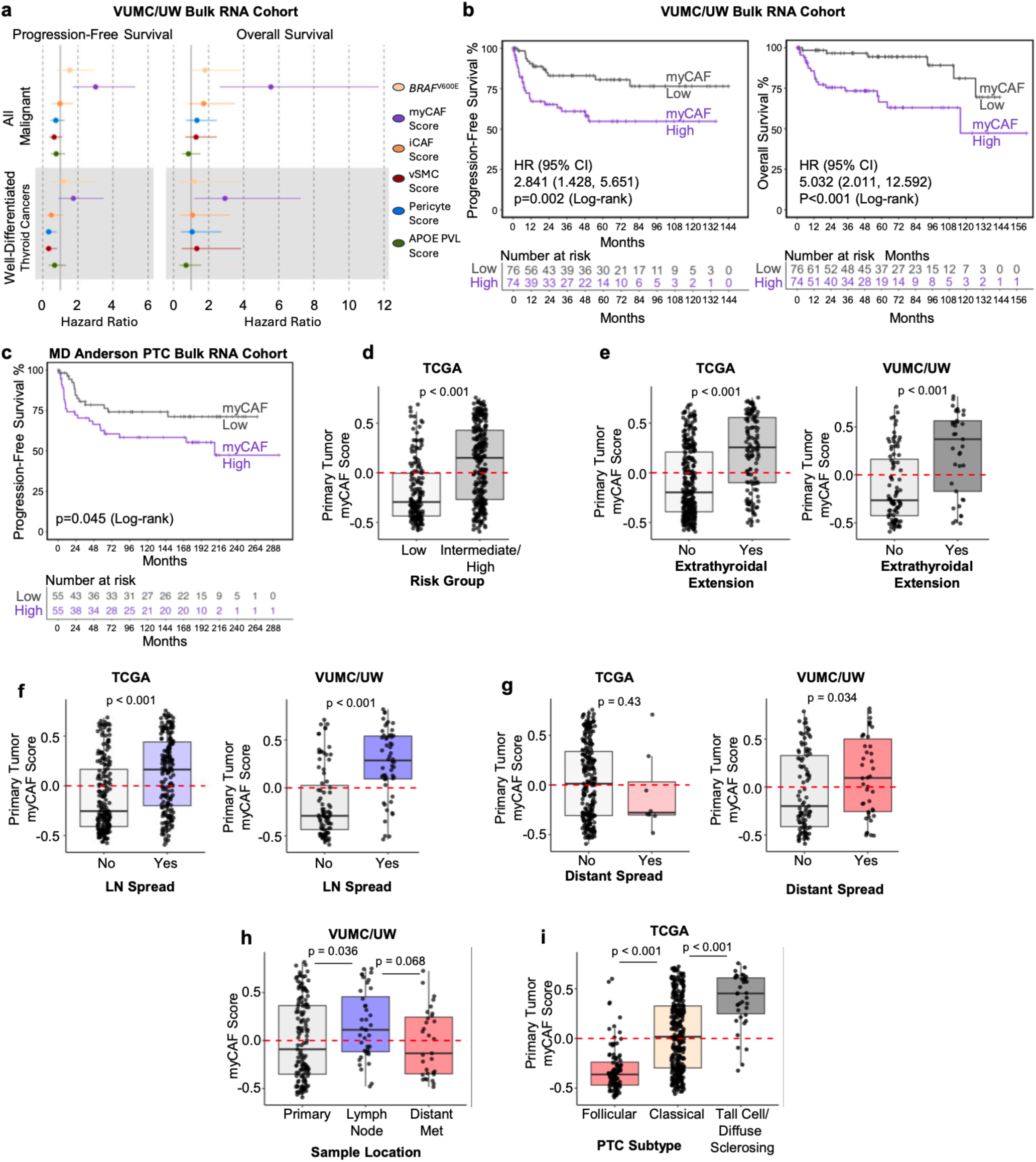
myCAFs are associated with aggressive thyroid cancer tumor phenotypes. a, Progression-free survival (left, PFS) and overall survival (right, OS) forest plots with interquartile range hazard ratios and 95% confidence intervals for the Vanderbilt University Medical Center/University of Washington thyroid cancer bulk RNA-sequencing cohort (VUMC/UW bulk RNA cohort). The cohort is split into all malignant samples (top) or well-differentiated thyroid cancers (bottom). Hazard ratios are shown for *BRAF*^V600E^ mutation and stromal cell single-sample gene set variation analysis scores (myofibroblast cancer-associated fibroblast score, myCAF score; inflammatory cancer-associated fibroblast score, iCAF score; APOE+ perivascular-like score, APOE+ PVL score; pericyte score; vascular smooth muscle cell score, vSMC) generated from gene lists from **Supplementary Table 2**. **b**, PFS (left) and OS (right) survival curves showing the VUMC/UW malignant cohort split into myCAF-high and myCAF-low by 50^th^ percentile myCAF score. **c**, PFS curve of MD Anderson papillary thyroid cancer (PTC) bulk RNA-sequencing cohort split into myCAF-high and myCAF-low by 50^th^ percentile myCAF score. P-values for **b-c** calculated with log-rank test. **d-g,** Boxplots depicting primary tumor myCAF score in (**d**) The Cancer Genome Atlas (TCGA) PTCs stratified by risk group as defined by the 2009 American Thyroid Association guidelines,^40^ (**e**) TCGA PTCs (left) and VUMC/UW malignant cohort (right) stratified by the presence or absence of extrathyroidal extension, (**f**) TCGA PTCs (left) and VUMC/UW malignant cohort (right) stratified by the presence or absence of lymph node spread, and (**g**), TCGA PTCs (left) and VUMC/UW malignant cohort (right) stratified by the presence or absence of distant metastasis. P-values calculated with Wilcoxon rank sum test. **h**, myCAF score in the VUMC/UW malignant cohort stratified by sample location. **i**, Primary tumor myCAF score TCGA PTCs stratified by histology subtype. Tall cell and diffuse sclerosing PTC subtypes are grouped. P-values for **h,i** calculated with Wilcoxon rank sum test with Bonferroni correction.

### myCAFs are enriched in epithelial ATCs but not mesenchymal ATCs

Our previous work utilizing MAP score initially identified two ATC subsets with unique stromal infiltration, high-MAP ATCs with robust CAF infiltrate and low-MAP ATCs with minimal CAF infiltrate.^25^ To further refine these subsets and investigate which ATCs are associated with myCAF infiltrate, we performed subclustering of the ATC tumor cell clusters in our single-cell atlas. ATC tumor cells largely clustered by sample, consistent with the heterogeneity in gene expression and copy number alterations that is a hallmark of ATC (Fig. 5a).^20,28^ However, distinguishing clusters by the presence or absence of *BRAF*^V600E^ mutation indicated that despite the heterogeneity of these tumors, *BRAF*^V600E^ and *BRAF* WT ATCs have distinct gene expression patterns (Fig. 5b). Through differential abundance and differential gene expression analysis, we identify epithelial ATCs, enriched for high-MAP tumors with *BRAF*^V600E^, and mesenchymal ATCs, enriched for low-MAP tumors and *BRAF* WT tumors. Specifically, we identified that epithelial ATCs are often *BRAF*^V600E^ mutant with a more epithelial, keratin-rich gene expression program and a robust CAF infiltrate. Rather than having robust CAF recruitment, mesenchymal ATCs are typically *BRAF* WT ATCs that demonstrate a highly mesenchymal gene program, expressing their own array of extracellular matrix genes (Fig 5c-f, Extended Data Fig. 4a-c). As identified by Lu et al., mesenchymal ATCs also express *CDK6.*^28^ Consistent with their mesenchymal gene expression, TGFβ signaling is activated in *BRAF* WT ATCs whereas MAPK signaling is activated in *BRAF*^V600E^ ATCs (Extended Data Fig. 4d). To confirm that these *BRAF* WT mesenchymal tumor cells are not fibroblasts, we performed copy number inference^41^ and observed *FAP*-expressing clusters with aneuploid tumor cells (Extended Data Fig. 4e,f). In contrast, *BRAF*^V600E^ tumors have diploid clusters of *FAP*-expressing CAFs (Extended Data Fig. 4g,h).

**Fig. 5:**
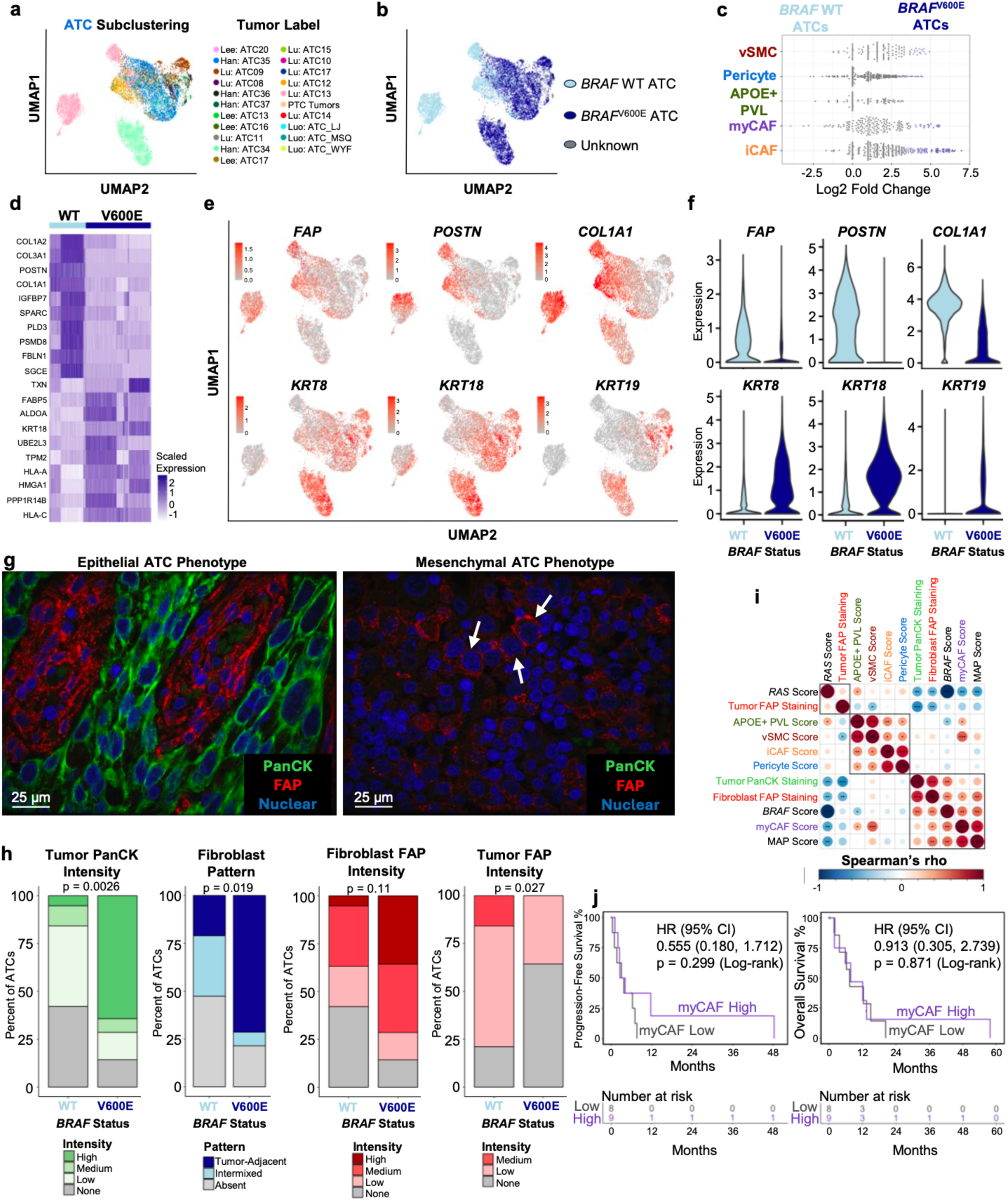
myCAFs are enriched in epithelial ATCs. a,b, Uniform Manifold Approximation and Projection (UMAP) showing subclustering of anaplastic thyroid carcinoma (ATC) tumor cells colored by (**a**) sample (tumor label) or (**b**) *BRAF*^V600E^ mutation status. **c**, Milo differential abundance testing of cancer-associated fibroblast (CAF) and perivascular stromal cell populations (inflammatory CAF, iCAF; myofibroblast CAF, myCAF; APOE+ perivascular-like, APOE+ PVL; pericyte; vascular smooth muscle cell, vSMC) between ATC tumors with and without *BRAF*^V600E^ mutations. Individual dots depict neighborhoods calculated by Milo. Coloring of individual neighborhoods as dark blue (*BRAF*^V600E^) or light blue (*BRAF* wild type, WT) indicates differential abundance with a spatial false discovery rate (FDR) of less than 0.1. Neighborhoods colored light grey have a spatial FDR greater than 0.1. **d**, Heatmap showing scaled expression of the top 10 differentially expressed genes with expression in at least 80 percent of cells in the population of interest between *BRAF* WT and *BRAF*^V600E^ ATCs. **e**, UMAP of ATC subclustering colored by expression of myCAF genes *FAP*, *POSTN*, and *COL1A1* (top row) or epithelial keratins *KRT8*, *KRT18*, and *KRT19* (bottom row). **f**, violin plots of showing expression of genes from **d** in ATC tumor cells split by *BRAF* status (WT vs V600E mutation). **g,** Representative multiplex immunofluorescence images from staining of 33 ATCs in the Vanderbilt University/University of Washington bulk RNA-sequencing cohort (VUMC/UW cohort) for pan-cytokeratin (PanCK, green) and FAP (red). Left: representative image of ATC with PanCK+ tumor cells and tumor-adjacent FAP+ fibroblasts. Right: representative image of ATC with no PanCK staining, FAP+ tumor cells, and minimal stromal FAP staining. White arrows point to malignant nuclei with membranous FAP staining. **h,** Bar plots showing pathologist scoring of ATC multiplex immunofluorescence. ATC tumors are split by *BRAF* status (WT vs V600E). P-values calculated with Fisher’s exact test. **i,** Corrplot showing Spearman’s rho correlations for the ATC multiplex immunofluorescence staining samples comparing PanCK tumor cell intensity, FAP tumor cell intensity, FAP fibroblast intensity, single-sample gene set variation analysis (ssGSVA) scores for stromal subpopulations, ssGSVA score for molecular aggression and prediction (MAP) score, *BRAF* score, and *RAS* score in the VUMC/UW ATC cohort. Axes ordered by hierarchical clustering. Boxes indicate hierarchical clustering groups. Significance levels indicate *p<0.05, **p<0.01, or ***p<0.001. **j,** Progression-free survival (left) and overall survival (right) survival curves showing VUMC/UW cohort ATCs split by 50^th^ percentile myCAF score. P-values calculated by log-rank test.

To morphologically visualize and confirm these differences, we performed multiplex immunofluorescence for pan-cytokeratin and FAP across 33 ATCs (19 *BRAF* WT, 14 *BRAF*^V600E^) from the VUMC/UW bulk RNA-sequencing cohort. We histologically confirmed these two distinct ATC patterns: 1) epithelial ATCs had pan-cytokeratin-expressing tumor cells with extensive FAP+ tumor-adjacent stroma (epithelial ATCs); and 2) mesenchymal ATCs had pan-cytokeratin negative tumor cells with frequent FAP- expression and minimal fibroblasts (mesenchymal ATCs) (Fig. 5g). Quantification indicated that *BRAF*^V600E^ tumors have greater tumor pan-cytokeratin intensity (p=0.0026), a tumor-adjacent fibroblast pattern (p=0.019), and a trend toward greater fibroblast FAP intensity (p=0.11). In contrast, *BRAF* WT tumors had greater intensity of FAP tumor staining (p=0.027), indicating that *BRAF*^V600E^ ATCs were predominantly “epithelial” whereas *BRAF* WT were “mesenchymal,” though the relationship was not mutually exclusive (fig. 5h).

We next correlated this staining data to MAP score and stromal GSVA scores. MAP score previously separated ATCs into CAF-rich (MAP-high) and CAF-depleted (MAP-low) tumors.^25^ We observe that ATC fibroblast FAP staining has a significant correlation with myCAF score, MAP score, and greater *BRAF*- like phenotype, but not other stromal populations. Pan-cytokeratin intensity had a significant correlation with *BRAF*-like phenotype (Fig. 5i). Although our survival analysis across bulk RNA-sequencing cohorts indicates that CAF-rich tumors have worse outcomes, that may not be the case within universally aggressive ATCs.^19^ When comparing PFS and OS between myCAF-high and myCAF-low ATCs, we observed no difference. (Fig. 5j). Importantly, this is a historic, retrospective ATC cohort and these patients did not receive targeted or immunotherapy. Overall, we have identified two distinct ATCs: an epithelial ATC, commonly *BRAF*^V600E^, with keratin expression and fibroblast infiltrate, and a mesenchymal ATC with few fibroblasts.

### Spatial mapping of CAFs in tumor progression

Based on the clinical relevance of CAFs in tumor progression, we investigated the spatial localization of CAF populations. We generated spatial transcriptomic data across the spectrum of disease progression, including eight *RET*-fusion driven pediatric PTCs, five *BRAF*^V600E^ PTCs, and 15 ATC samples, seven of which spatially capture WDTC as it de-differentiates into ATC (Extended Data Fig. 5a,b). In *BRAF* WT ATCs, we observed ubiquitous *CDK6* expression and heterogeneous expression of CAF markers in tumor cell regions, consistent with tumor cell expression of a CAF-like mesenchymal profile (Extended Data Fig. 5c-e).

Because our spatial transcriptomics data contains multiple cells for each spatial barcode, we used our single-cell atlas to dissect the localization of individual cell types as a percent of each spatial barcode. Spatial mapping of tumor cell and CAF subpopulations revealed a striking pattern of tumor-adjacent myCAFs and tumor-distant iCAFs across pediatric PTCs, adult PTCs, and ATCs (Fig. 6a). We quantified the minimum distance of myCAFs and iCAFs from tumor cells in PTCs and *BRAF*^V600E^ ATC samples containing both myCAFs and iCAFs (n = 18). iCAFs were further from tumor cells across samples as compared to myCAFs (p=0.0047, Fig. 6b,c). Immunohistochemistry for the myCAF protein POSTN and the iCAF protein APOD confirmed the tumor-adjacent and tumor-distant localization (Fig. 6d). In one pediatric PTC with background Hashimoto thyroiditis, we observe mapping of iCAFs to surrounding areas of lymphocytic thyroiditis, supporting fibroblast infiltrates with iCAF-like inflammatory phenotypes in thyroid autoimmune disease (Extended Data Fig. 6a).

**Fig. 6:**
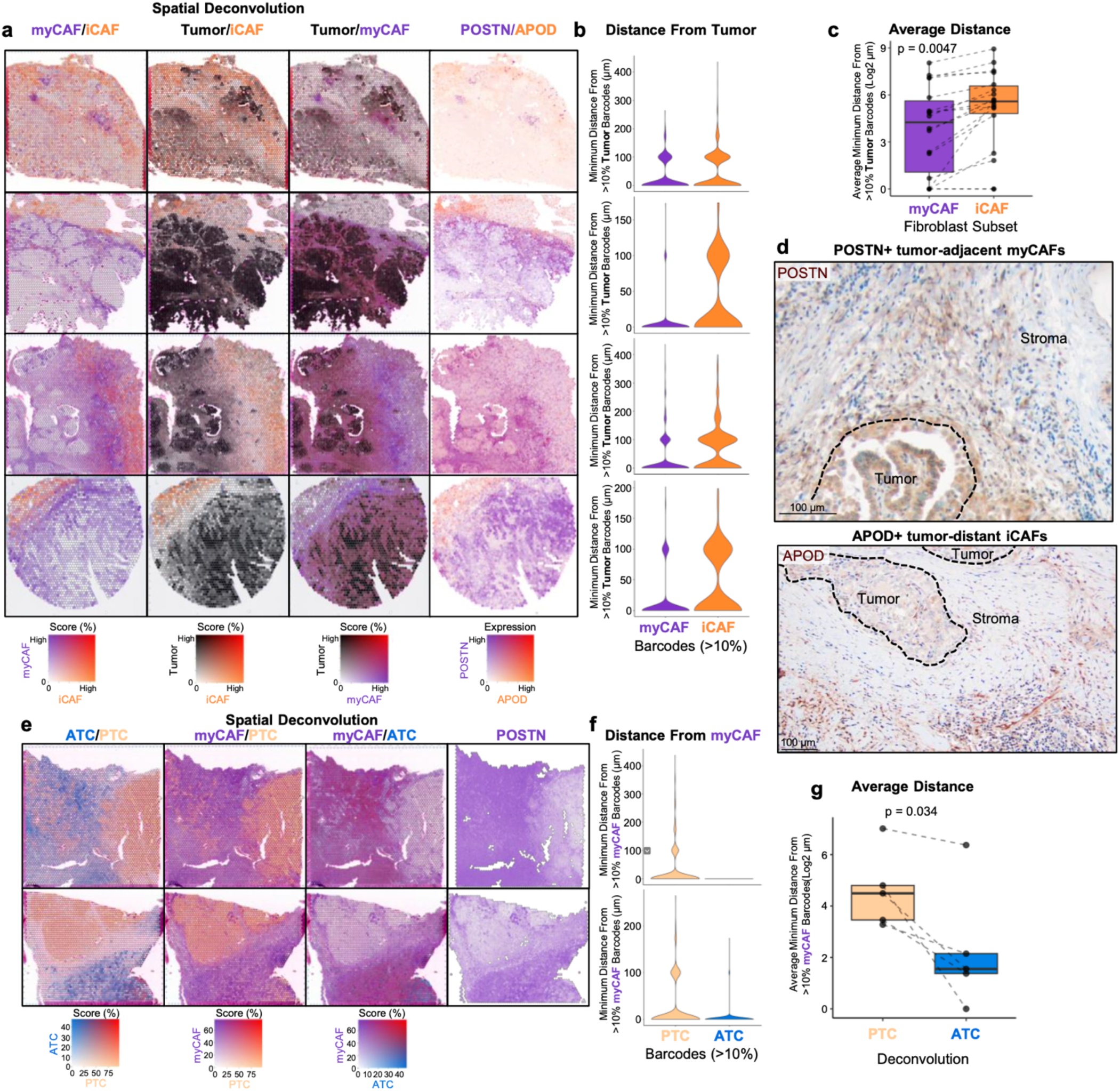
Spatial localization of CAF populations in thyroid cancer progression. a, Spatial feature plots from left to right of myofibroblast cancer-associated fibroblast (myCAF, purple), inflammatory CAF (iCAF, orange), and tumor cell (black) Robust Cell Type Decomposition (RCTD) and myCAF (*POSTN*, purple) and iCAF (*APOD*, orange) marker gene expression. Tumor cell deconvolution is the sum of papillary thyroid cancer (PTC) and anaplastic thyroid cancer (ATC) deconvolution. From top to bottom samples shown are a pediatric PTC (Peds04), adult PTC (Thy7), adult PTC with associated ATC (Thy5), and an adult ATC (Thy4). Mixing of deconvoluted populations or marker genes is shown by a color gradient that becomes red. **b,** Violin plots pertaining to samples from **a** showing the minimum Euclidean distance from a spatial barcode with at least 10% tumor RCTD for each spatial barcode with at least 10% myCAF RCTD (purple) or iCAF RCTD (orange). **c**, Boxplot showing the average of spatial barcode minimal Euclidean distances from **b** for myCAF (purple) and iCAF (orange) across 18 spatial transcriptomics samples with *BRAF*^V600E^ mutations and the presence of both myCAF and iCAF RCTD populations. P-value calculated with paired t-test. **d**, Representative images of immunohistochemistry for POSTN (top) and APOD (bottom) at the invasive edge of PTC. **e**, Spatial feature plots of two mixed PTC/ATC samples (Thy6, Thy11) showing ATC (blue), PTC (light orange), and myCAF (purple) RCTD or myCAF (*POSTN*, purple) marker gene expression. Mixing of deconvoluted populations is shown by a color gradient that becomes red. **f**, Violin plots pertaining to samples from **e** showing the minimum Euclidean distance from a spatial barcode with at least 10% myCAF RCTD for each spatial barcode with at least 10% PTC RCTD (light orange) or ATC RCTD (blue). **g,** Boxplot showing the average of spatial barcode minimal Euclidean distances from **f** for PTC (light orange) and ATC (blue) across 5 mixed PTC/ATC tumors with *BRAF*^V600E^ mutations. P-value calculated with paired t-test.

To comprehensively understand the stromal picture of thyroid tumors, we explored the spatial localization of myCAFs, pericytes, and vSMC. For evaluation of myCAFs, we assessed five *BRAF*^V600E^ samples with PTC tumor cells de-differentiating into ATC. We observed abundant myCAFs with POSTN expression infiltrating ATC regions, but only lining the periphery of PTC regions (Fig. 6e, Extended Data Fig. 6b). Distance quantification indicated closer proximity of ATC cells to myCAFs across all five tumors (p=0.034, Fig. 6f,g), quantitatively confirming increased myCAF infiltration as tumors progress from PTC to ATC. Because pericytes were upregulated in PTC single-cell samples, we next evaluated PTC specimens with adjacent stroma (Fig. 7a, Extended Data Fig. 6c). As expected, myCAFs mapped to the invasive perimeter of PTC areas. In contrast to myCAFs, pericytes mapped within central areas of PTC (Fig. 7b,c, Extended Data Fig. 6d,e). This association with pericytes was quantitatively confirmed using spatial cross-correlation analysis across 12 PTCs with mapping of both pericyte and myCAF populations (p=0.024, Fig. 7d-f, Extended Data Fig. 6f). Pathologist annotation of PTC regions demonstrated pericyte deconvolution was concentrated within the papillary vascular cores of PTC (Fig. 7g-h, p=0.029). Immunohistochemistry further confirmed RGS5+ pericytes within the papillary fibrovascular core of PTCs (Fig. 7i). vSMCs localized to large perivascular areas (pathologist annotated) on five tumors (Fig. 7j-m, Extended Data Fig. 6g-i). Immunohistochemistry confirmed αSMA+ vSMCs lining large vessels within tumors (Fig. 7n). In conclusion, we spatially mapped key stromal cell populations in thyroid cancer and identified a robust increase in tumor-adjacent myCAFs with de-differentiation in *BRAF*^V600E^ tumors.

**Fig. 7:**
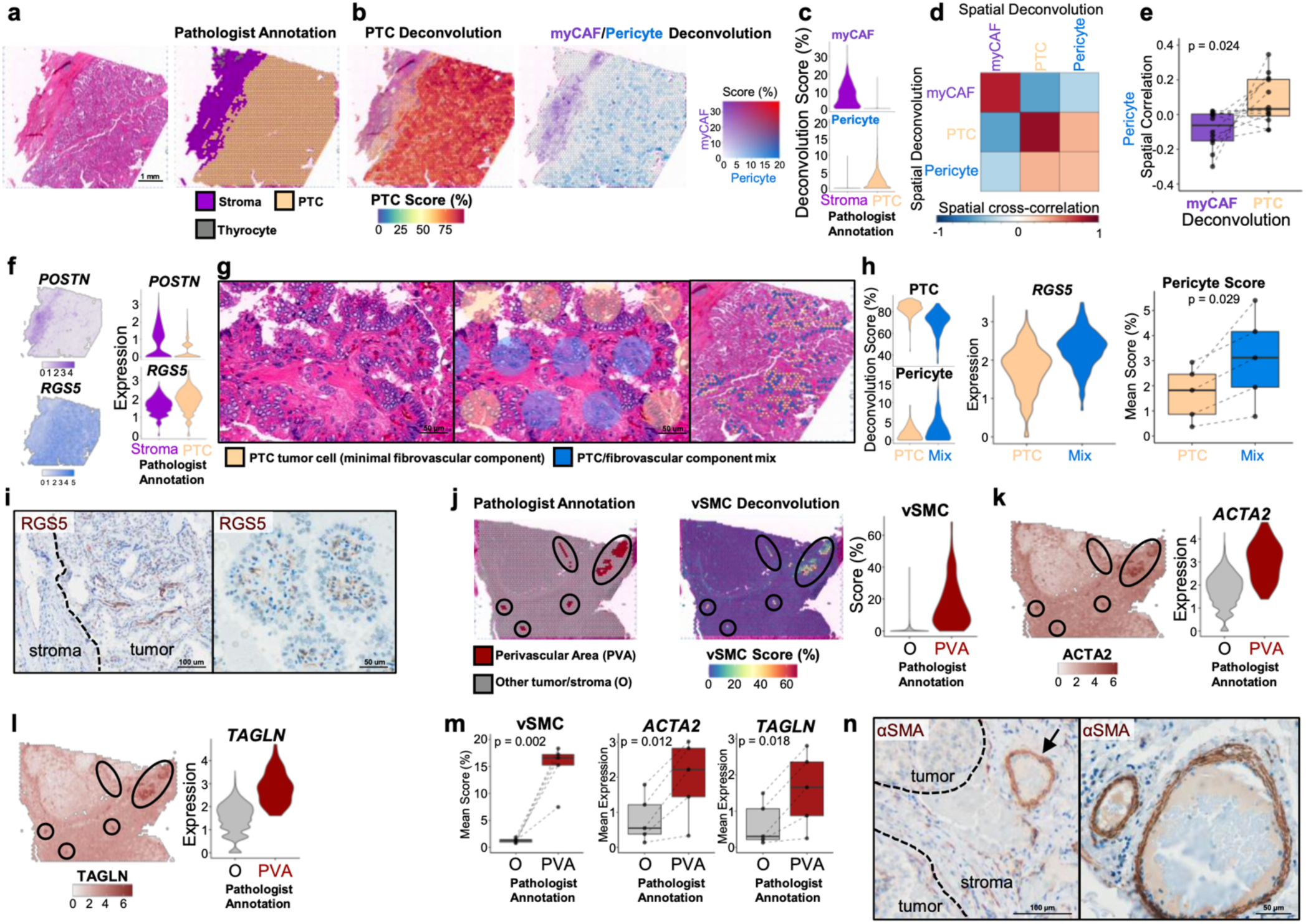
Spatial localization of perivascular populations in thyroid cancer. a, Hematoxylin and eosin staining of representative papillary thyroid cancer (PTC) Thy15 (left) with pathologist annotation of PTC (light orange) and stromal (purple) spatial barcodes (right). **b**, Spatial feature plots of representative PTC Thy15 showing Robust Cell Deconvolution (RCTD) of PTC (left, red color gradient) and myofibroblast cancer-associated fibroblast (myCAF)/pericyte (right, purple/blue color gradients) populations. **c**, Violin plots showing myCAF and pericyte RCTD scores within pathologist annotated stromal (purple) and PTC (light orange) barcodes. **d,** Correlation heatmap showing MERINGUE spatial cross-correlation of myCAF, PTC, and pericyte RCTD scores (ordered by hierarchical clustering) for representative PTC Thy15. **e,** Boxplot showing pericyte spatial cross-correlation with myCAF (purple) and PTC (light orange) across 12 PTC samples containing PTC and stromal regions. P-value calculated with paired t-test. **f**, Spatial feature plots (left) and violin plots (right) depicting myCAF (*POSTN*) and pericyte (*RGS5*) marker gene expression in pathologist annotated stromal and PTC regions. **g**, Pathologist annotation of PTC tumor regions into predominantly PTC tumor cell (light orange) or a mix of PTC tumor cells and fibrovascular cells (blue). Left: zoomed in hematoxylin and eosin staining. Middle: zoomed in hematoxylin and eosin staining with spatial barcodes labeled as PTC or PTC/fibrovascular component mix. Right: zoomed out hematoxylin and eosin staining with a sample of the spatial barcodes labeled as PTC or PTC/fibrovascular component mix. **h,** Violin plots showing PTC and pericyte RCTD deconvolution scores (left) and pericyte marker *RGS5* expression (middle) across spatial barcodes labeled as PTC (light orange) or PTC/fibrovascular component mix (mix). Right: box plots depicting quantification of pericyte RCTD deconvolution scores in PTC (light orange) or PTC/fibrovascular component mix (blue) regions across five samples (Thy15, Thy16, Thy17, Peds07, Peds08). P-value calculated with paired t-test. **i**, RGS5 immunohistochemistry of representative PTC. **j**, Pathologist spatial annotation of large perivascular areas (PVAs) and spatial feature plot of vascular smooth muscle cell (vSMC) RCTD scores. Left: Pathologist annotation. Middle: vSMC RCTD score plot. Right: violin plot of vSMC RCTD score stratified by barcodes whether spatial barcodes are in large PVAs. **k-l**, Spatial feature plots (left) and associated violin plots (right) for vSMC marker genes (**k**) *ACTA2* and (**l**) *TAGLN*. **m**, Boxplots showing average vSMC RCTD score (left) or vSMC marker gene expression (right) in large PVA areas verses non-PVA tumor areas for five tumors with pathologist annotation (**supplemental** Fig. 6g). P-values calculated with paired t-test. **n,** Immunohistochemistry of αSMA in a representative PTC. Black arrow highlights a large perivascular area.

### myCAFs abut invasive tumor cells

CAFs are known to modulate tumor cell behavior to support cellular invasion. Our group recently discovered a CAF-adjacent leading-edge tumor cell population in *BRAF*^V600E^ ATC and invasive PTC with strong expression of *TNC* and striking phenotypic resemblance to a LAMB3+TNC+ partial epithelial-mesenchymal transition (pEMT) phenotype.^43,44^ Thus, we investigated a potential interaction between invasive pEMT cells and myCAFs in PTC and ATC. Subclustering PTC cells identified a population with enrichment of the pEMT signature from Puram et al. and increased interactions with myCAFs (Fig. 8a-c, Extended Data Fig. 7a,b, p=0.015). Spatial mapping of pEMT-PTC cells across pediatric and adult PTCs, as well as PTCs de-differentiating into ATCs, highlighted a *LAMB3*+ pEMT invasive cellular population directly abutting myCAFs, which we further confirmed with immunohistochemistry for LAMB3 (Fig. 8d-g). We performed spatial ligand-receptor interaction analysis across four PTCs with robust pEMT-PTC populations. All four showed increased interaction between pEMT-PTC cells and myCAFs relative to non- pEMT-PTC cells (p=0.13), with top upregulated signaling pathways including collagen, laminin, and tenascin (Fig. 8h,i, Extended Data Fig. 7c-e).

**Fig. 8:**
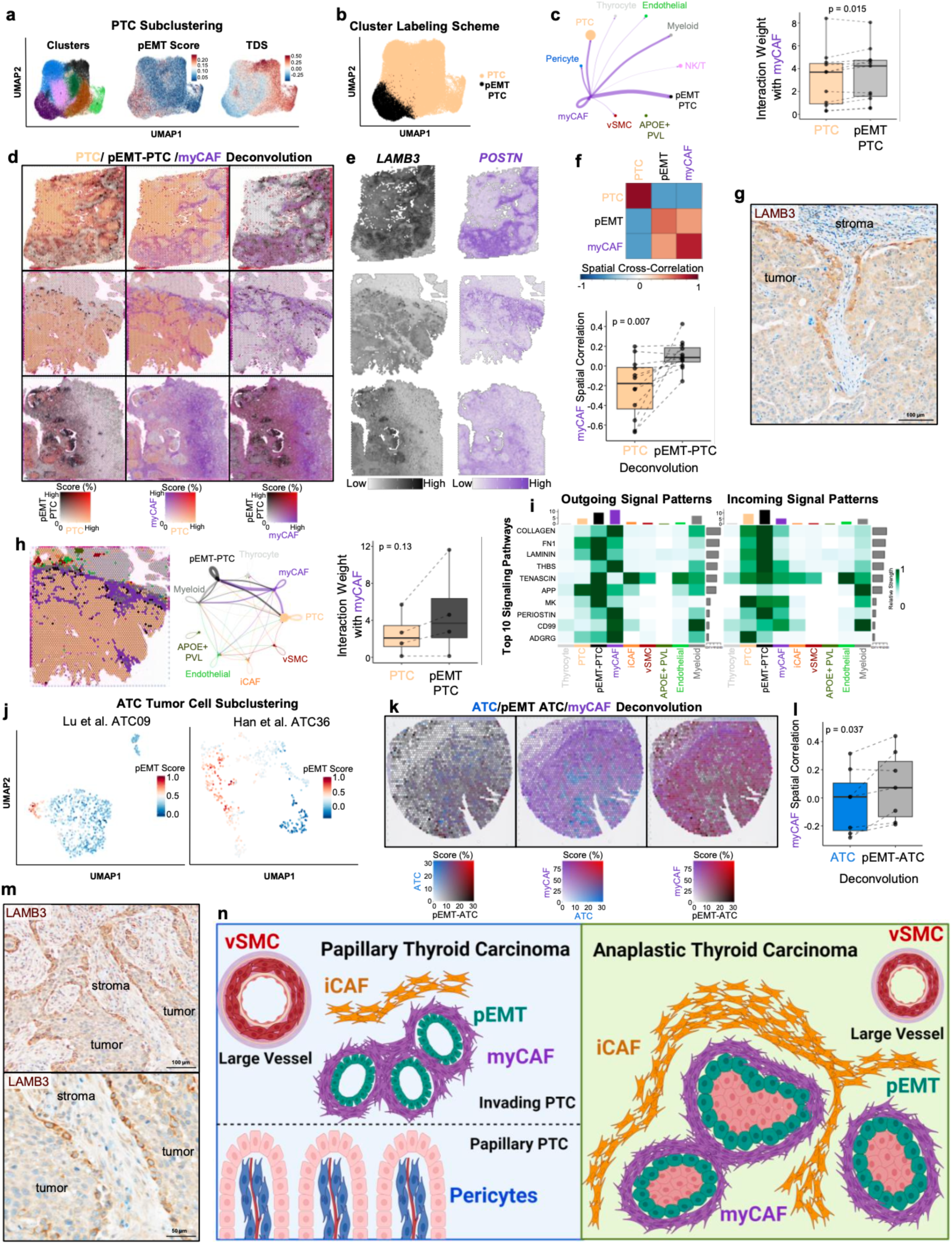
Spatial mapping reveals a partial EMT phenotype in invasive tumor cells. a, Uniform Manifold Approximation and Projection (UMAP) of thyroid cancer atlas papillary thyroid cancer (PTC) cluster subclustering colored by clusters (left), partial epithelial-mesenchymal transition (pEMT) score (middle), or thyroid differentiation score (TDS, right). **b,** UMAP showing labeling of PTC subclusters as either PTC (light orange) or pEMT-PTC (black). **c,** Interaction weights between myofibroblast cancer-associated fibroblasts (myCAFs) and other single-cell populations in PTC samples. Left: Interaction weights between myCAFs and other cell clusters in representative PTC WJL from Luo et al. The width of purple lines depict interaction weights. Right: Box plots depicting quantification of interaction weights between myCAFs and PTC (light orange) or pEMT-PTC (grey) in 11 PTC single-cell samples with myCAF, PTC, and pEMT- PTC populations. P-value calculated with paired t-test. **d-e,** Spatial feature plots of (**d**) PTC (light orange), pEMT-PTC (black), and myCAF (purple) Robust Cell Type Decomposition (RCTD) scores or (**e**) pEMT- PTC (*LAMB3*, black) and myCAF (*POSTN*, purple) marker gene expression. Mixing of deconvoluted populations or marker genes is shown by a color gradient that becomes red. From top to bottom samples shown are a pediatric PTC (Peds08), an adult PTC (Thy7), and an adult mixed PTC/ATC (Thy5). **f**, Spatial cross-correlation analysis between myCAF, PTC, and pEMT-PTC populations. Top: correlation heatmap showing MERINGUE spatial cross-correlation of PTC, pEMT-PTC, and myCAF RCTD scores ordered by hierarchical clustering for representative PTC Peds08. Bottom: boxplot showing myCAF spatial cross-correlation with PTC (light orange) and pEMT-PTC (grey) across 12 PTC samples containing PTC and pEMT-PTC phenotypes. P-value calculated with paired t-test. **g,** Immunohistochemistry of LAMB3 at the leading-edge of a representative PTC. **h,** Ligand-receptor interaction weights in PTC with quantification across four samples. Left: labeling of spatial barcodes by the population with the highest RCTD score in representative PTC Thy7. Middle: spatial ligand-receptor interaction weights between labeled populations on the left. The width of lines indicates interaction weights. Right: Boxplot showing myCAF interaction weights with PTC (light orange) and pEMT-PTC (grey) across four PTCs that had sufficient pEMT-PTC spatial barcodes for ligand-receptor interaction analysis. P-value calculated with Wilcoxon signed-rank test. **i**, Heatmap showing the top 10 signaling patterns in representative PTC Thy7 with outgoing signal scores on the left and incoming signal scores on the right. The bottom depicts the labeled spatial barcode populations. **j,** UMAP of ATC tumor cell subclustering colored by Seurat module scores for Puram et al. pEMT genes for Lu et al. ATC09 (left) and Han et al. ATC36 (right). **k,** Spatial feature plots of Thy4 with RCTD scores generated by replacing tumor cell clusters with Lu et al. ATC09 subclustering. Left: pEMT-ATC RCTD (black) and ATC RCTD (blue). Middle: ATC RCTD (blue) and myCAF RCTD (purple). Right: pEMT-ATC RCTD (black) and myCAF RCTD (purple). Mixing of deconvoluted populations or marker genes is shown by a color gradient that becomes red. **l,** Boxplot showing myCAF spatial cross-correlation with ATC (blue) and pEMT-ATC (grey) across 7 *BRAF*^V600E^ ATC samples with pEMT and myCAF populations. P-value calculated with paired t-test. **m,** Immunohistochemistry of LAMB3 in a representative *BRAF*^V600E^ mutant ATC. **n**, Schematic overview of fibroblast populations across thyroid cancer progression. Abbreviations: iCAF, inflammatory cancer-associated fibroblast; vSMC, vascular smooth muscle cell; PVL, perivascular-like; NK/T, natural killer/T cell

Because ATC tumors subcluster by sample, we looked for pEMT populations within individual *BRAF*^V600E^ tumor samples. In two ATCs we identified a subset of tumor cells with enrichment of pEMT genes and increased interaction with myCAFs (Fig. 8j, Extended Data Fig. 7f,g). Spatial mapping of ATC and pEMT- ATC tumor cells highlighted a pEMT leading-edge phenotype intermixed with myCAFs in *BRAF*^V600E^ ATCs (Fig. 8k,l). Immunohistochemistry for LAMB3 further confirmed a myCAF-adjacent leading-edge pEMT phenotype (Fig. 8m). The pEMT gene signature used to identify leading edge cells was associated with aggressive tumor phenotypes across bulk RNA-sequencing cohorts (Extended Data Fig. 8a-h). Altogether, we show tumor-adjacent myCAFs directly communicating with invasive, pEMT-like tumor cells in both aggressive PTCs and ATCs (Fig. 8n).

## Discussion

Given the mounting evidence that CAFs shape tumor behavior and immunity across cancers, there is interest in therapeutic targeting of CAFs for treatment of highly-lethal ATC.^25,27,39,45–49^ However, pan-fibroblast targeting strategies are unlikely to be effective,^4–6^ and specific CAF subset modulation in thyroid cancer is hindered by the lack of comprehensive profiling of subtypes and spatial localization. To discover CAF heterogeneity and function in thyroid cancer, we created a thyroid cancer single-cell sequencing atlas^26–32^ and spatially resolved CAF dynamics in invasive tumors at sites of de-differentiation from papillary to anaplastic morphology. Through this work, we identify CAF subsets that serve as critical mediators of cancer progression, inflammatory thyroiditis, and tumor hemodynamics. myCAFs in thyroid cancer are centrally located partners in invasion, directly abutting and communicating with invading tumor cells to alter extracellular matrix and differentiation programs. In contrast, iCAFs are tumor-distant, enriched in areas of lymphocytic thyroiditis, supporting recent work showing fibroblasts with CAF phenotypes in non-neoplastic conditions.^36^ We also distinguish a population of vSMCs that regulate large vessel hemodynamics in thyroid tumors. Finally, we show that pericytes localize within the tumor fibrovascular cores, likely regulating hemodynamics locally within tumor papillae. Additional research is needed to explore the potential of pericytes to serve as CAF precursors, given the similarities in gene expression between myCAFs and pericytes in thyroid cancer.

Spatial profiling of CAF-tumor interactions in thyroid cancer has numerous implications for patient care. Using multiple large patient cohorts across institutions, we confirm that myCAFs are associated with disease aggression and worse outcomes in thyroid cancer. Clinical diagnostic assays to detect myCAFs in thyroid cancer could potentially identify high-risk patients and improve thyroid cancer prognostication. Additionally, targeting the direct interactions between myCAFs and invading tumor cells may represent a promising therapeutic strategy for invasive disease. Targeting of iCAFs in thyroid cancer may help to increase patient response to immunotherapy. For patients with inflammatory autoimmune thyroid conditions, the modulation of iCAFs may serve to reduce thyroid destruction and improve functionality. Targeting of pericyte populations in thyroid cancer could represent a promising strategy for altering tumor hemodynamics and reducing tumor growth and vascularization. Finally, an understanding of the stromal organization of ATC based on our proposed classification as either epithelial or mesenchymal may allow for improved and personalized therapeutic decision making in this highly lethal disease.

The primary limitation of our study is the resolution of the Visium sequencing platform without matched single-cell sequencing. While our single-cell atlas mitigates this limitation, additional large studies with paired samples are needed to better spatially characterize epithelial and mesenchymal ATCs. A second limitation is that ATC single-cell samples have limited numbers of tumor cells due to necrosis and low tumor purity. Finally, recent studies have indicated that *BRAF*^V600E^ ATCs may have better response to immunotherapy combined with targeted therapies than *BRAF* WT ATCs.^50^ Because our cohorts are retrospective and did not receive immunotherapy, we are not able to assess the impact of ATC subtyping into epithelial and mesenchymal categories on the response to immunotherapy and targeted therapy. Future work is needed to understand how targeting the robust stroma in epithelial ATCs impacts their response to therapy.

In conclusion, our findings elucidate tumor-CAF interactions spatially across thyroid cancer progression using a combination of single-cell, spatial, and bulk RNA-sequencing cohorts. We identify CAF populations important for thyroid physiology, with myCAFs specifically serving as critical modulators of invasion in thyroid cancer. The dynamic changes in CAFs as thyroid cancer de-differentiates have broad implications across solid tumors, where the identification of tumor-supporting tumor-stroma interactions could revolutionize understanding of tumor biology and lead to novel CAF-targeting strategies.

## Methods

### Generation of thyroid cancer single-cell atlas

#### Sample pre-processing and individual analysis

Raw count matrices and meta data were downloaded from seven published thyroid cancer single-cell RNA-sequencing data sets (Gene Expression Omnibus accession codes: GSE184362, GSE193581, GSE232237, GSE182416, GSE191288; Genome Sequence Archive accession number: HRA000686; CNGB Nucleotide Sequence Archive of China National GeneBank accession number: CNP0004262).^26–32^ R package Seurat 4.4.0 was used throughout pre-processing and atlas integration.^34^ For each sample, Seurat objects were generated and pre-processed to exclude cells with fewer than 200 unique features, fewer than 500 total counts, or greater than 10 percent mitochondrial reads. Doublet detection was run using R package scDblFinder 1.14.0 with default arguments and predicted doublets were removed.^51^ Sample NORM21 from Lu et al. was excluded from further analysis due to less than 200 cells passing quality control and sample PTC10_T from Pu et al. was excluded due to concern over ambient RNA limiting clear separation of tumor and stromal clusters on independent sample analysis. Following pre- processing, each sample was normalized and variance stabilized using Seurat function SCTransform with vst.flavor set to “v2.”^52,53^ Normalized counts were annotated with human primary cell atlas (HPCA) labels using SingleR.^54^ Copy number inference was performed using R package CopyKAT1.1.0 using default settings with normalized counts as input to identify aneuploid cells.^41^ Samples were individually clustered and annotated with broad cell type labels (malignant, non-malignant thyrocyte, stromal, myeloid, NKT, endothelial, and B/plasma cell) based on HPCA labels, presence or absence of aneuploid cells, provided cell type labels from each sample’s reference paper when available, and expression of canonical marker genes. Broad cell type and HPCA labels were added to sample meta data prior to integration.

#### Sample integration and broad cell type labeling

Samples were integrated using fast mutual nearest neighbors (fastMNN) within R package SeuratWrappers 0.2.0.^55^ First, raw RNA counts and meta data were merged from all individual samples. To limit over batch correcting biologically distinct cell types, samples were merged in order of heterogeneity of cell populations. Tumors containing PTC cells, ATC cells, and diverse microenvironment populations were merged first and tumors with few microenvironment populations were merged later.

The merged data was normalized and integrated with SeuratWrappers command RunFastMNN based on the top 3000 variable features. The top 30 dimensions within the MNN integration were used for Seurat commands RunUMAP, FindNeighbors, and FindClusters. Broad clusters were determined at a resolution of 0.6 and visualized with Uniform Manifold Approximation and Projection (UMAP). Broad clusters were labeled based on consensus of HPCA labels, canonical marker genes, provided cell type labels from reference papers, and cell type meta data from individual sample analysis. Because the stromal and ATC clusters overlapped, each cell within this group was assigned as stromal or ATC based on the meta data from individual sample analysis. Within analysis of individual samples, stromal clusters and ATC clusters were distinct and differentiated by aneuploidy profiles. The identity of thyrocyte and tumor cell populations was further validated by generating module scores in Seurat with published TCGA PTC gene lists for thyroid differentiation, *BRAF*-like thyroid tumors, and *RAS*-like tumors, and with a 12 gene panel for distinguishing ATC from PTC published by Han et al.^29,33^

#### Stromal subclustering

Stromal cell labels were extracted from the broad labels of the integrated thyroid atlas and used to subset raw RNA counts and meta data from the merged atlas. The subset RNA count data were integrated and clustered as described above in sample integration at a resolution of 0.2. Differential gene expression between each stromal subcluster and all other stromal cells was calculated using Seurat function FindMarkers with the test specified as “MAST.”^56^ Results were plotted with Seurat function DoHeatmap for the top 15 genes from each subcluster with expression in at least 50 percent of cells based on descending fold-change. Genes with log2 fold-change of at least one were included as marker genes for each cluster (Supplementary Table 2) and used for gene ontology analysis. Additionally, pathway activity scores were generated for each stromal cell using R package PROGENy 1.22.0^57,58^ and summarized across stromal subclusters following the analysis pipeline from Figure 3 of Gao et al. (https://github.com/aliceygao/pan-Fibroblast).^7^ The identity of stromal subclusters was informed by generating module scores in Seurat with published CAF subtype and perivascular gene lists.^8–15,35,36^ Module scores were plotted using Seurat function DotPlot. Module scores and marker genes were also mapped onto the stromal UMAP using Seurat function FeaturePlot.

#### Tumor cell subclustering

ATC and PTC cell labels were independently extracted from broad labels of the integrated thyroid atlas and used to subset raw RNA counts and meta data from the merged atlas. The subset RNA count data were integrated and clustered as described above in sample integration at a resolution of 0.3. For ATC subclustering, differential gene expression analysis was performed between *BRAF* WT and *BRAF*^V600E^ cells. The top 10 marker genes with at least 80% expression in the population of interest were plotted with DoHeatmap in order of descending fold-change. For PTC subclustering, a pEMT module score was generated using a published pEMT gene signature from head and neck squamous cell carcinoma.^44^ PTC subclusters were labeled as “pEMT-PTC” or “PTC” based on their enrichment of the pEMT module score.

#### Differential abundance testing

Differential abundance testing of stromal subclusters and epithelial clusters was performed with R package MiloR 1.8.1 (https://github.com/MarioniLab/miloR).^59^ For buildGraph and makeNhoods functions, parameters k and d were both set to 30 and 10 percent of cells were sampled for makeNhoods. Results were plotted with function plotDAbeeswarm. Neighborhoods with a spatial FDR of less than 0.1 were colored corresponding to the condition they were enriched in.

#### Ligand-receptor interaction analysis

Ligand-receptor interaction analysis was performed for individual samples within the integrated atlas with R package CellChat 2.1.2 using published pipelines.^60,61^ Communication probabilities were calculated using triMean. Interaction weights between myCAFs and tumor cell populations were compared for samples containing at least 100 pEMT-PTC and 100 non-pEMT-PTC tumor cells. Interaction weights between cell populations were plotted with CellChat function netVisual_circle. The top 10 signaling pathways were plotted with CellChat function netAnalysis_signalingRole_heatmap.

### Bulk RNA-sequencing analysis

#### Sequencing cohorts

Stromal and tumor cell subpopulation abundance and disease associations were analyzed in four thyroid cancer bulk RNA-sequencing cohorts: the VUMC/UW cohort (312 samples spanning benign and malignant neoplasms) TCGA cohort (496 PTCs), CNUH/SNUH cohort (263 normal thyroids, 348 PTCs, 5 PDTCs, 16 ATCs), and MD Anderson cohort (110 PTCs).^31,33^ Sequencing and processing of the VUMC/UW cohort has been previously described.^25^ For this study, raw counts were Transcripts Per Million (TPM) normalized. TCGA sequencing, meta, and clinical data was downloaded from cBioPortal For Cancer Genomics (cbioportal.org), including previously calculated *BRAF*-*RAS* scores, American Thyroid Association risk stratification based on the 2009 guidelines, and whether tumors had lymph node and/or distant metastases.^33,40,62,63^ TCGA PTC fibrosis scores from previously published imaging analysis were associated with stromal subpopulation abundance.^39^ The CNUH/SNUH cohort was downloaded from Gene Expression Omnibus (GSE213647) and TPM normalized. The 110 PTCs in the MD Anderson cohort were processed and sequenced using previously described methods.^64^ Raw counts were TPM normalized for this study.

### Calculating gene expression scores

Gene expression scores for all bulk RNA-sequencing cohorts were generated via single-sample gene set variation analysis (GSVA) with R package GSVA 1.48.3 using default arguments.^65^ GSVA scores were generated for each stromal subcluster marker gene list (Supplementary Table 2), Puram et al. published pEMT marker genes, and our previously published 549 gene thyroid cancer MAP score.^25,44^ For the MD Anderson, VUMC/UW, and CNUH/SNUH cohorts, TPM normalized counts were used as input. For the TCGA cohort, RSEM normalized counts were used as input. BRS was previously calculated for TCGA and VUMC/UW cohorts.^25,33^

### Survival analysis

110 PTCs from the MD Anderson and 150 malignant samples from the VUMC/UW cohort had clinical follow-up data for progression-free survival (PFS) and overall survival (OS) analysis. These cohorts were split into two groups by 50^th^ percentile for stromal and tumor subpopulation scores of interest and survival analysis was performed as previously described.^25^ Kaplan-Meier survival curves were compared and tested using the log-rank test. Cox proportional hazards models were also calculated for stromal and tumor subpopulation scores and summarized with forest plots. In the forest plots, the hazard ratio with a 95% confidence interval was reported for an interquartile range (IQR) increase when scores were analyzed as continuous variables.

### Spatial transcriptomics analysis

#### Platform

The Visium Spatial Gene Expression v1 for formalin-fixed paraffin embedded (FFPE) tissue platform was used to generate spatial transcriptomics data (10x Genomics, Pleasanton, CA).

#### FFPE block selection and sequencing

Eight thyroid carcinoma formalin fixed paraffin embedded (FFPE) blocks with ATC histology were previously processed by our lab (Extended Data Fig. 5 samples Thy2-4, Thy8, Thy10, Thy14, Thy19 and Thy20).^25^ An additional 20 FFPE blocks, including 8 pediatric PTCs, 5 adult PTCs, and 7 ATCs were processed. Tumor regions were chosen by pathologist review (VLW) to spatially capture areas of progression from PTC to ATC and areas with mixed tumor and stroma. One of the previously processed ATCs (Thy10) had mixed PTC and ATC histology and six of the newly processed ATCs contain PTC components.

#### Sample processing and sequencing

5 µm FFPE tissue sections were cut onto Visium Gene Expression Slides (Visium Spatial Gene Expression Slide Kit, PN-1000188). Slides were subsequently stained, imaged, processed, and sequenced as previously described.^25^ The Visium Human Transcriptome Probe Set v1.0 was used and sequencing was performed using the NovaSeq 6000 platform (Illumina, San Diego, CA).

#### Data processing

Visium sequencing FASTQ files and 20X image scans were pre-processed with Space Ranger 2.0.0 (10x Genomics, Pleasanton, CA). Space Ranger outputs were loaded into R package Seurat 4.4.0 using function Load10X_Spatial or R package SingleCellExperiment 1.22.0 using function readVisium.^34,66^ Subsequent analyses were performed with either Seurat objects or SingleCellExperiment objects depending on the compatibility of the published analysis method. Spatial sequencing reads were normalized and variance stabilized using Seurat function SCTransform with vst.flavor set to “v2.”^52,53^

#### Enhanced resolution gene expression plots

Individual genes were plotted at subspot resolution using BayesSpace 1.10.1 following the pipeline in the package vignette (https://github.com/edward130603/BayesSpace).^67^

#### Cell type deconvolution of spatial barcodes

Raw RNA counts and broad cluster labels from the integrated thyroid cancer single-cell RNA-sequencing atlas was used as a reference for deconvolution of spatial transcriptomics data. The broad stromal cluster labels were split into iCAF, myCAF, APOE+ PVL, pericyte, and vSMC subpopulations based on stromal subclustering labels. Deconvolution was repeated with the PTC cluster split into PTC and pEMT-PTC groups based on PTC subclustering results and with the tumor cells subset to only ATC09 from Lu et al. with ATC and pEMT-ATC distinction. Deconvolution was performed with R package spacexr 2.2.1 using the RCTD algorithm following the pipeline for Visium spatial transcriptomics samples (https://github.com/dmcable/spacexr).^68^ The run.RCTD function was run with doublet_mode set to “full” to accommodate the resolution of Visium spatial transcriptomics data. Determining malignant cell fraction of spatial barcodes in *BRAF* wild type ATCs was performed with R package SpaCET 1.2.0 (https://github.com/data2intelligence/SpaCET) with cancer type set to “PANCAN.”^42^

#### Annotation of spatial barcodes

Spatial barcodes were annotated using Loupe Browser 8.1.1 (10x Genomics, Pleasanton, CA) by a practicing pathologist with greater than 10 years of experience (VLW) and a pathologist trainee. In five samples (Peds02, Thy5, Thy11, Thy9, and Thy13), barcodes were assigned as within either a large perivascular area or tumor/other area. Barcodes from two representative PTCs (Thy15 and Thy17) were annotated as tumor or stroma. Within PTC regions of five PTCs (Thy15, Thy16, Thy17, Peds07, and Peds08), barcodes were further split into pure PTC barcodes and barcodes containing a mix of PTC tumor cells and fibrovascular cells found at the central core of PTCs. Pathologist annotations were used to validate the localization of stromal cell deconvolution with RCTD.

#### Distance calculations

For each sample, Visium barcode coordinates were extracted from Seurat objects using the Seurat command GetTissueCoordinates with scale set to NULL. To convert the coordinates from pixels to µm, the coordinates of each sample were multiplied by 65 (the approximate diameter in µm of each Visium spatial barcode) and divided by the full resolution spot diameter in pixels (the full resolution spot diameter is generated by Space Ranger and located in the scalefactors_json.json file). Using the tissue coordinates in µm, the Euclidean distance was calculated between individual coordinates. A distance matrix was generated for each sample containing sequencing barcodes as both rows and columns with values containing the distance between barcode pairs. For minimum distance calculations, distance matrices were subset to rows and columns containing spatial barcodes of interest. For example, the minimum distance for barcodes with at least 10% PTC normalized RCTD score weight from barcodes with at least 10% myCAF normalized RCTD score weight was calculated by subsetting the rows of the distance matrix to PTC barcodes and the columns to myCAF barcodes and returning the minimum distance value in each row. For analysis across samples, the average minimum distance was calculated for each sample of interest.

#### Spatial cross-correlation

Spatial cross-correlation analysis between RCTD scores for single-cell populations was performed with R package MERINGUE 1.0 (https://github.com/JEFworks-Lab/MERINGUE).^69^ Visium barcode coordinates were extracted from Seurat objects using the Seurat command GetTissueCoordinates with default arguments. Spatial neighbors were calculated by inputting barcode coordinates to the MERINGUE function getSpatialNeighbors with filter distance set to 7.5. Spatial neighbors and a matrix of normalized RCTD score weights were input into MERINGUE function spatialCrossCorMatrix to calculate spatial cross-correlation statistics between deconvoluted cell populations. The resulting spatial cross-correlation matrix was plotted with R package corrplot 0.92 using the corrplot function with order set to “hclust” to group populations with similar spatial localization. Boxplots of spatial cross-correlation between populations of interest were plotted across samples using ggplot2 3.5.0.^70^

#### Spatial ligand-receptor interaction analysis

Spatial ligand-receptor interaction analysis was performed with R package CellChat 2.1.2 using the v2 human database subset to protein signaling cell-cell communication genes and pathways. Spatial barcodes identities for ligand-receptor interaction analysis were assigned based on the cell type composing a majority of the barcode following RCTD. SCT transformed expression data were used as input. Communication probabilities were computed using triMean, an interaction range of 250 µm, a contact dependent interaction range of 100 µm, and a minimum number of Visium barcodes per group of 10, as recommended in CellChat tutorials (https://github.com/jinworks/CellChat).^60,61^

### Tissue staining of FFPE tissue

#### Multiplex immunofluorescence

Multiplex immunofluorescence staining of 33 ATC FFPE blocks (19 *BRAF* WT, 14 *BRAF*^V600E^) for fibroblast activating protein alpha (Abcam ab207178 recombinant rabbit monoclonal anti-fibroblast activation protein alpha (FAP) IgG, clone EPR20021, 1:100, Abcam, Cambridge, UK) and pan-cytokeratin (eBioscience 53-9003-82 mouse monoclonal anti-pan cytokeratin IgG1 AF488, clone AE1/AE3, 1:100, Thermo Fisher, Waltham, MA) was performed as previously described.^25^ Representative multiplex immunofluorescence images were scored by a practicing pathologist with greater than 10 years of experience and thyroid expertise (VLW). For each ATC tissue section, FAP staining of stromal cells was categorized as none, low, medium, or high, and FAP staining of tumor cells was categorized as none, low, or medium. Tumor cells were identified by nuclear morphology. An overall fibroblast pattern was assigned based on FAP staining of stromal cells as absent, intermixed, or tumor adjacent. Tumor cell pan-cytokeratin was scored as none, low, medium, or high. The proportions for these staining categories were compared between *BRAF* wild type and *BRAF*^V600E^ ATCs using Fisher’s exact test and plotted as bar plots using R package ggplot2 3.5.0.^70^ The pancytokeratin and FAP staining scores were given rank order from least to highest staining and correlated with bulk RNA GSVA scores generated from the same tumors using Spearman’s rank correlation coefficient. Correlations were calculated and plotted using R package corrplot 0.92.

#### Immunohistochemistry

5 µm tissue sections were cut from FFPE blocks of PTC and ATC. Deparaffinization and antigen retrieval were performed as previously described for multiplex immunofluorescence staining of FFPE.^25^ After antigen retrieval, tissues were treated with BLOXALL endogenous blocking solution, peroxidase and alkaline phosphatase (SP-6000-100, Vector Laboratories, Newark, CA) for 10 min, washed with 0.05% Tween 20 in PBS, and blocked for 2 h with 10% goat serum in PBS (blocking buffer). Primary antibodies (Abcam ab5694 rabbit polyclonal anti-alpha smooth muscle actin IgG, 1:500; Abcam ab314670 rabbit recombinant monoclonal anti-RGS5 IgG, clone EPR28539-64, 1:500; Invitrogen PA5-21514 rabbit polyclonal anti-laminin beta-3 IgG, 1:100; Invitrogen PA5-34641 rabbit polyclonal anti-periostin IgG, 1:100; Sigma-Aldrich HPA040520 affinity isolated rabbit polyclonal anti-APOD IgG, 1:500) were diluted in blocking buffer and incubated on tissue sections at 4°C for 16 h (Abcam, Cambridge, UK; Thermo Fisher, Waltham, MA; MilliporeSigma, Burlington, MA). Tissue sections were washed with 0.05% Tween 20 in PBS and incubated for 30 minutes with ImmPRESS HRP horse anti-rabbit IgG polymer (MP-7801- 15, Vector Laboratories, Newark, CA). Tissue sections were washed with 0.05% Tween 20 in PBS and developed via incubation for 2 min with an equal mix of reagent 1 and reagent 2 of ImmPACT DAB EqV

Substrate (MP-7801-15, Vector Laboratories, Newark, CA). Following development, tissue sections were rinsed in distilled water, counterstained for 3 min with Mayer’s Hematoxylin (MHS32-1L, MilliporeSigma, Burlington, MA), rinsed in tap water, washed for 1 min in Scott’s water (10 g magnesium sulfate, 2g sodium bicarbonate, 1 L water), rinsed in tap water, dehydrated (1 min each of 25% ethanol, 50% Ethanol, 70% ethanol, 100% ethanol, and 3 min xylene), and cover slipped.

### Data availability

The spatial transcriptomics data is available at the Human Tumor Atlas Network (HTAN) Data Portal (https://data.humantumoratlas.org/). Single-cell RNA-sequencing data sets were downloaded from publicly available databases (Gene Expression Omnibus accession codes: GSE184362, GSE193581, GSE232237, GSE182416, GSE191288; Genome Sequence Archive accession number: HRA000686; CNGB Nucleotide Sequence Archive of China National GeneBank accession number: CNP0004262). TCGA bulk RNA-sequencing data was downloaded from cBioPortal For Cancer Genomics (cbioportal.org). Bulk RNA-sequencing data from the CNUH/SNUH cohort was downloaded from Gene Expression Omnibus (accession number: GSE213647). The VUMC/UW bulk RNA-sequencing data set has restrictions on data sharing, as previously described.^25^ Aggregate-level data will be shared by lead contact (Dr. Vivian Weiss) upon request. Individual-level data is only available through collaboration following approval of the lead contact and The Vanderbilt University Medical Center IRB. The MD Anderson bulk RNA-sequencing data will be made available upon publication.

### Code availability

All code for reproducing analyses in the manuscript has been uploaded to GitHub in the following repository: https://github.com/mloberg16/THCA_Fibroblast_Atlas.

## Acknowledgements

Overview Fig. 8n was created using BioRender.com with relevant licenses obtained. This research was made possible through the American Association for Cancer Research. We thank the Vanderbilt Translational Pathology Shared Resource (TPSR) for their support in tissue sectioning and staining, the Vanderbilt Digital Histology Shared Resource (DHSR) for their support with slide scanning, the Vanderbilt Technologies for Advanced Genomics (VANTAGE) core facility for their support with sequencing spatial transcriptomics specimens, and the Vanderbilt technologies for advanced Genomics Analysis and Research Design for their support with processing spatial transcriptomics data using Space Ranger (10x Genomics, Pleasanton, CA). We thank the Vanderbilt Cell Imaging Shared Resources (CISR) core for providing support for confocal microscopy (5P30 CA68485-19, S10 OD023475-01A1, DK20593, DK58404, DK59637, and EY08126). This work was funded by the American Society of Cytopathology (Young Investigator Award to V.L.W), American Thyroid Association (2019-0000000090 to V.L.W.), NIH (R35GM122516, R01CA244188, and R01CA272875 to E.L.; VCORCDP K12CA090625, K08CA240901, and R01CA272875 to V.L.W.; F30CA281125 to M.A.L.; NIGMS T32GM007347 to the Vanderbilt MSTP supported M.A.L. and M.L.T.; U01CA294527 and R01DK103831 to K.S.L.), V Foundation for Cancer Research (Scholar Award to V.L.W.), Children’s Cancer Research Fund (Research Award to V.L.W.), and American Cancer Society (133934-CSDG-19-216-01-TBG and RSG-22-084-01-MM to V.L.W.).

---

**Extended Data Fig. 1.**
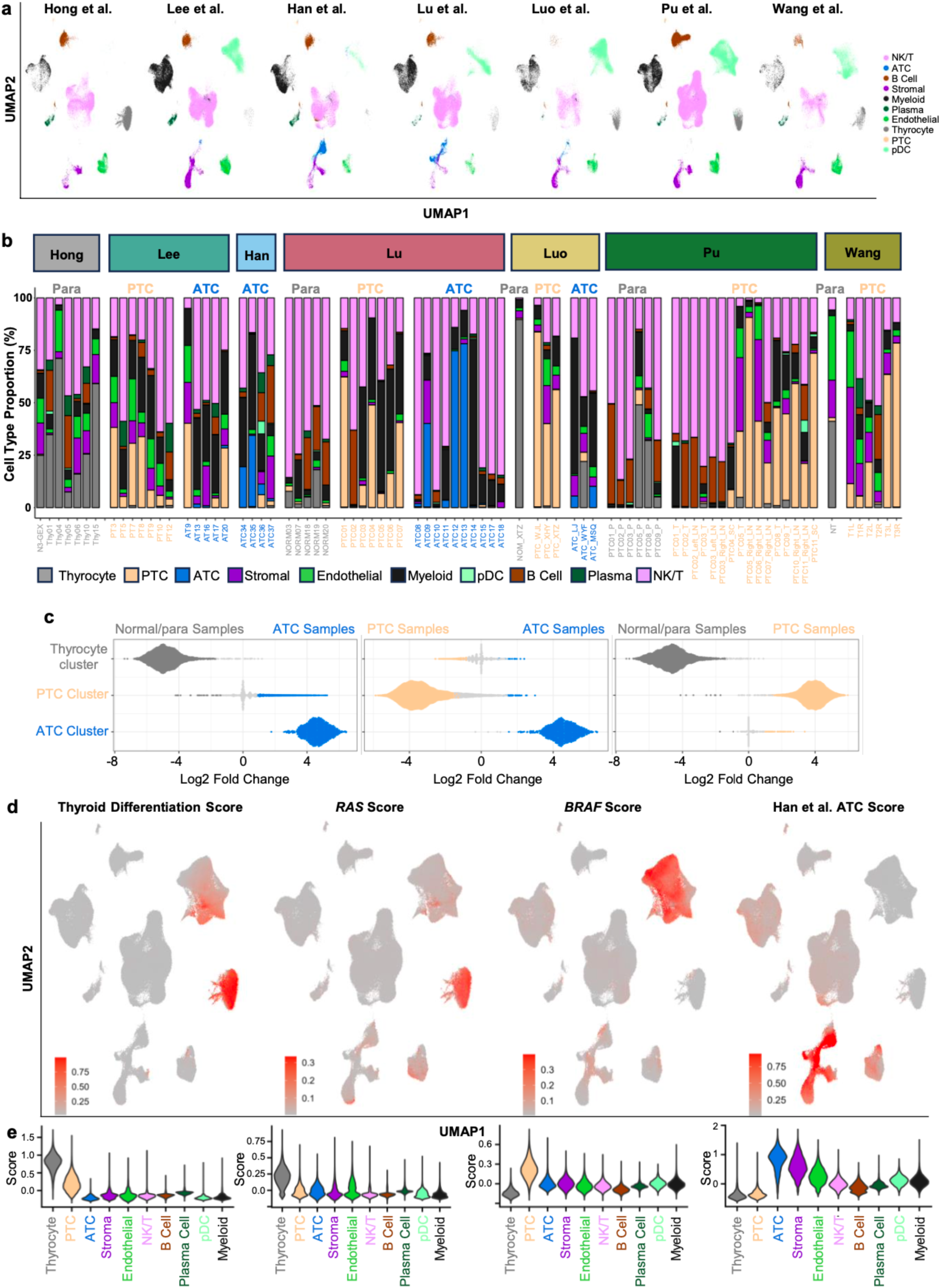
Integrated single-cell atlas validation of tumor cell populations. a, Broad cell population Uniform Manifold Approximation and Projection (UMAP) from Fig. 1b split by paper. **b**, Bar plots showing cell type proportion for each sample in the integrated thyroid cancer atlas. Papers and histologic subtype (anaplastic thyroid cancer, ATC; papillary thyroid cancer, PTC; normal/paratumor, Para) are labeled on top and sample names on bottom. **c**, Milo differential abundance testing of broad epithelial clusters (thyrocyte, PTC, ATC) between normal/paratumor samples and ATC samples (left), PTC samples and ATC samples (middle), and normal/paratumor samples and PTC samples (right). Individual dots depict neighborhoods calculated by Milo. Coloring of individual neighborhoods as dark grey (normal/paratumor), light orange (PTC), or blue (ATC) indicates a spatial false discovery rate (FDR) of less than 0.1. Neighborhoods colored light grey have a spatial FDR greater than 0.1. **d**, UMAP plot of integrated single-cell atlas showing module scores for thyroid differentiation, *RAS*-like genes, *BRAF*-like genes, and a 12 gene ATC score from Han et al. (left to right).^29,33^ **e**, Violin plots for the module scores in **d** depicting scores by broad cluster identification. Abbreviations: pDC, plasmacytoid dendritic cell; NK/T, natural killer/T cell.

**Extended Data Fig. 2:**
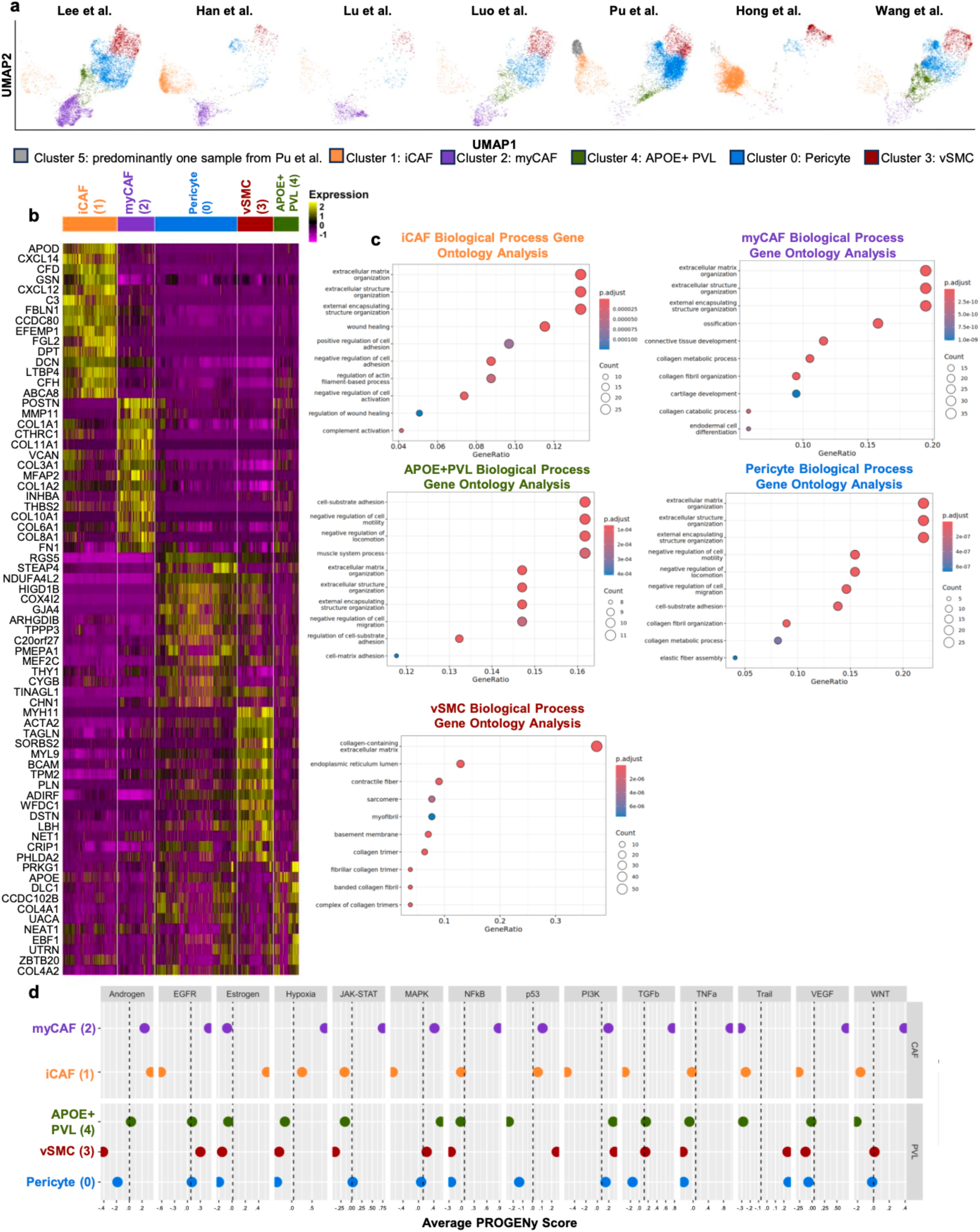
Enriched genes and pathways in stromal subclusters. a, Stromal cell subclustering Uniform Manifold Approximation and Projection (UMAP) from Fig. 2a split by paper. **b,** Heatmap showing scaled expression of the top 15 marker genes for each stromal subcluster with expression in at least 50 percent of cells in the cluster of interest. **c**, Gene ontology analysis of the genes for each stromal subcluster with log2 fold-change enrichment of at least 1.0 (**Supplementary Table 2** genes) showing the top 10 biological processes for each cluster. **d**, PROGENy pathway activity scores for each stromal subcluster.

**Extended Data Fig. 3:**
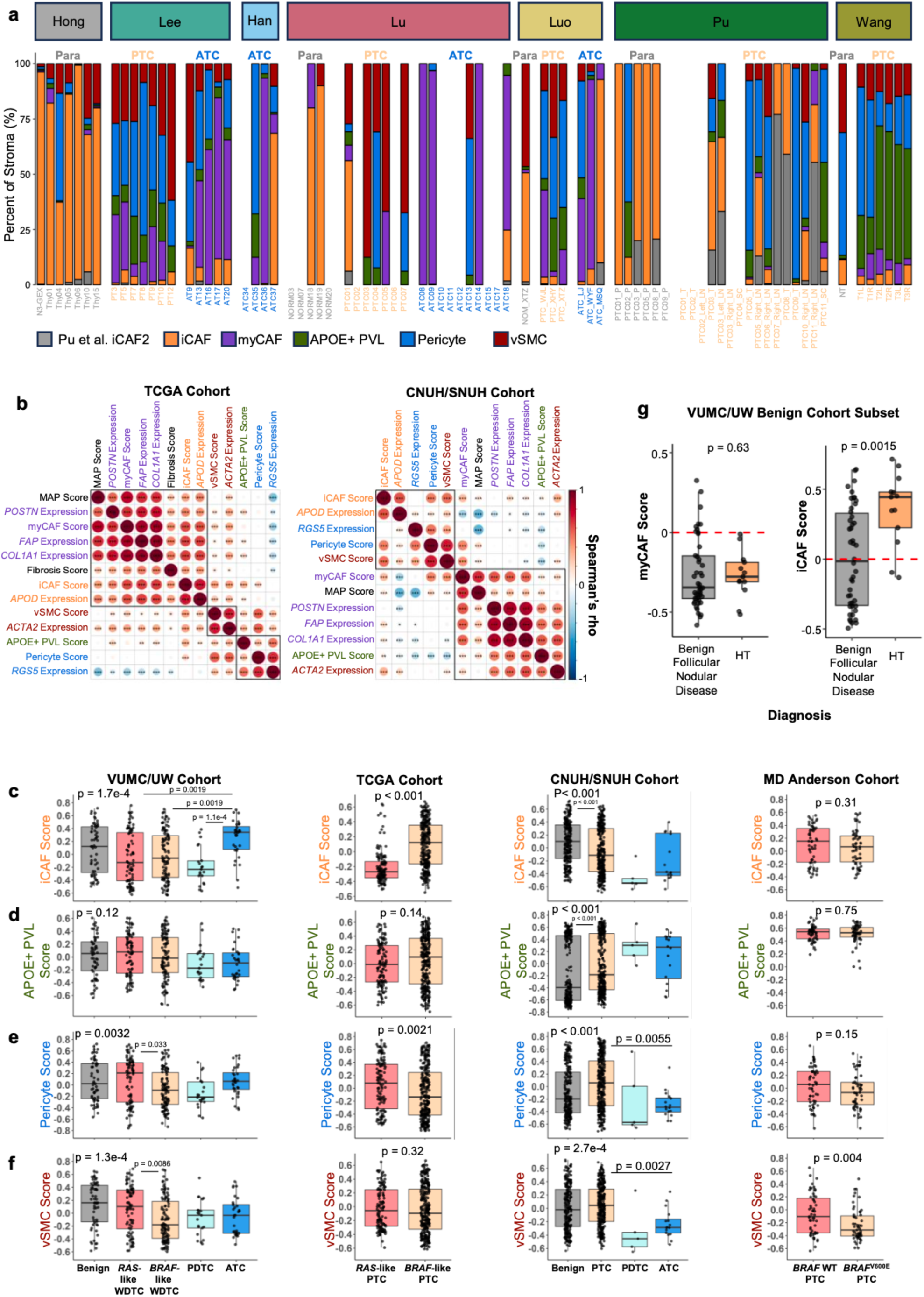
CAF and PVL stromal cell abundance in bulk and single-cell sequencing cohorts. a, Bar plots showing composition of stromal cells in integrated thyroid cancer single-cell atlas for each individual sample. Samples with a blank space had no stromal cells identified. **b,** Corrplot showing Spearman’s rho correlations between single-sample gene set variation analysis (ssGSVA) scores for stromal subpopulations, ssGSVA score for molecular aggression and prediction (MAP) score, and expression of marker genes for stromal subpopulations in The Cancer Genome Atlas PTC cohort (TCGA cohort, left) and Chungnam/Seoul National University Hospitals cohort (CNUH/SNUH cohort, right). Axes ordered by hierarchical clustering. Boxes indicate hierarchical clustering groups. Significance levels indicate *p<0.05, **p<0.01, or ***p<0.001. **c**-**f**, Boxplots showing ssGSVA scores for (**c**) inflammatory cancer-associated fibroblasts (iCAFs), (**d**) APOE+ perivascular-like cells (APOE+ PVL), (**e**) pericytes, and (**f**) vascular smooth muscle cells (vSMCs) by diagnosis across four distinct bulk RNA- sequencing cohorts. **g,** Boxplots showing myCAF ssGSVA scores (left) and iCAF ssGSVA scores (right) for benign thyroids split by diagnosis of benign follicular nodular disease or Hashimoto thyroiditis (HT). P-values for **c-g** calculated with Wilcoxon rank-sum test. Kruskal-Wallis test with subsequent pairwise Wilcoxon rank-sum tests with Bonferroni’s correction was used when comparing more than two groups. Abbreviations: myCAF, myofibroblast cancer-associated fibroblast.

**Extended Data Fig. 4:**
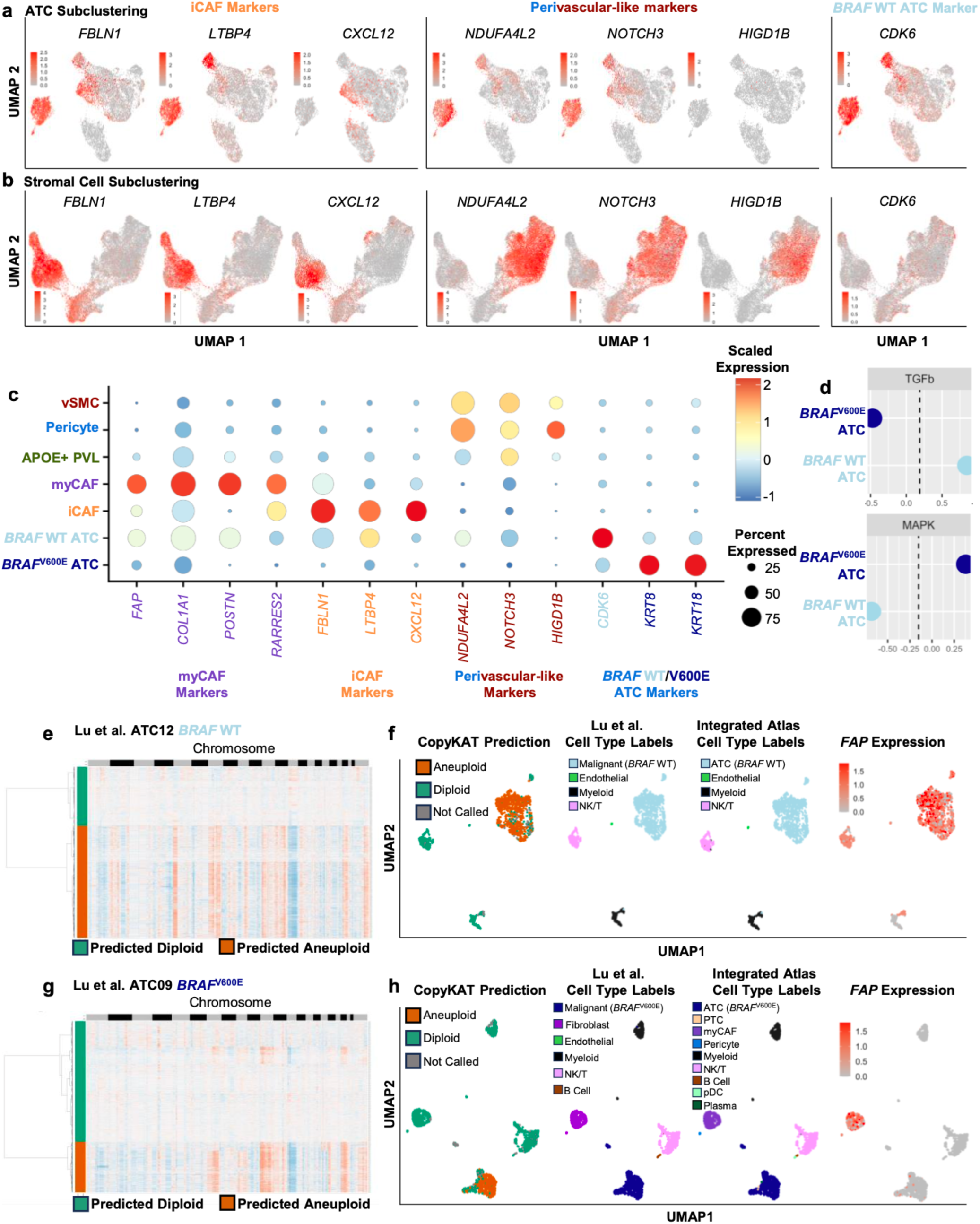
*BRAF* WT ATCs have heterogeneous expression of mesenchymal markers. a,b, Uniform Manifold Approximation and Projection (UMAP) plots of (**a**) anaplastic thyroid carcinoma (ATC) tumor cell subclustering and (**b**) stromal cell subclustering colored by expression of inflammatory cancer-associated fibroblast (iCAF) markers *FBLN1*, *LTBP4*, *CXCL12* (left), perivascular-like (PVL) markers *NDUFA4L2*, *NOTCH3*, *HIGD1B* (middle), and the ATC marker *CDK6* (right). **c**, Dot plot showing scaled expression of myofibroblast CAF (myCAF), iCAF, PVL, and ATC marker genes for stromal cell subpopulations (iCAF; myCAF; APOE+ PVL; pericyte; vascular smooth muscle cell, vSMC), *BRAF* wild type (WT) ATC tumor cells, and *BRAF*^V600E^ ATC tumor cells. **d**, PROGENy signaling pathway activity scores for *BRAF* WT and *BRAF*^V600E^ ATC cells. **e**, Heatmap showing gene expression (orange = high; blue = low) across chromosomal location (x-axis) for all cells (y-axis, cells in order of hierarchical clustering) from a representative *BRAF* WT ATC (ATC12 from Lu et al.) with CopyKAT aneuploid prediction shown (green = diploid; orange = aneuploid). **f**, UMAP of independent analysis of single-cell RNA-sequencing from ATC12 from Lu et al. colored from left to right by CopyKAT prediction, cell type labels provided by Lu et al., cell type labels from integrated thyroid cancer atlas, and *FAP* expression. **g**, Heatmap showing gene expression (orange = high; blue = low) across chromosomal location (x-axis) for all cells (y-axis, cells in order of hierarchical clustering) from a representative *BRAF*^V600E^ ATC (ATC09 from Lu et al.) with CopyKAT aneuploid prediction shown (green = diploid; orange = aneuploid). **h**, UMAP of independent analysis of single-cell RNA-sequencing from ATC09 from Lu et al. colored from left to right by CopyKAT prediction, cell type labels provided by Lu et al., cell type labels from integrated thyroid cancer atlas, and *FAP* expression.

**Extended Data Fig. 5:**
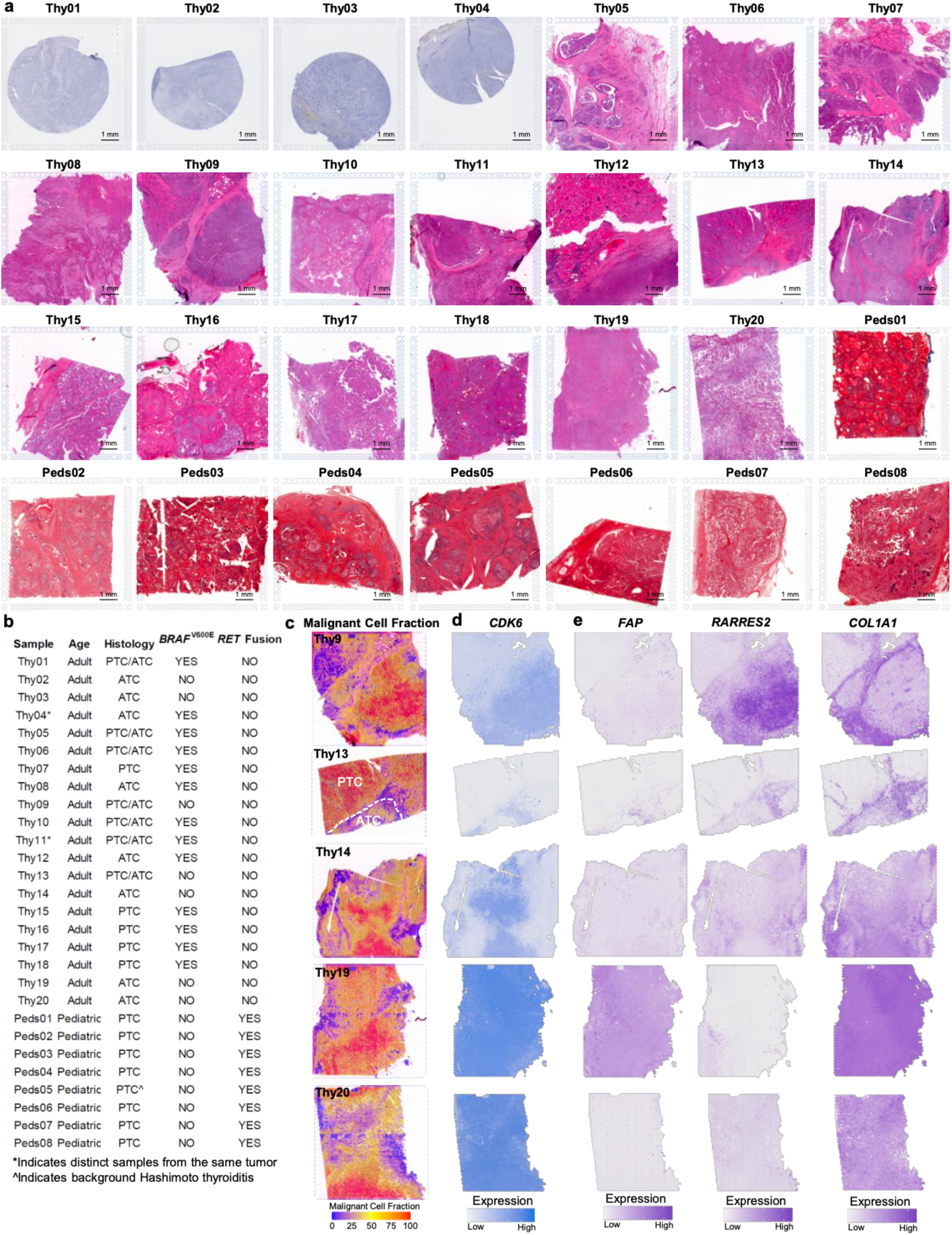
Thyroid cancer spatial sequencing atlas. a, Hematoxylin and eosin staining of 28 spatial transcriptomics tumor samples. **b,** Table with age, histologic subtype, and mutation data for 28 spatial transcriptomics tumor samples. *Indicates distinct samples from the same tumor. ^Indicates background Hashimoto thyroiditis. **c**, Malignant cell fraction of spatial transcriptomics barcodes for five *BRAF* wild type (WT) anaplastic thyroid carcinoma (ATC) tumors calculated by Spatial Cellular Estimator for Tumors (SpaCET).^42^ **d**, Expression of ATC marker *CDK6* across five *BRAF* WT ATC spatial transcriptomic samples. **e**, Expression of myCAF markers *FAP*, *RARRES2*, *COL1A1* across five *BRAF* WT ATC spatial transcriptomic samples.

**Extended Data Fig. 6:**
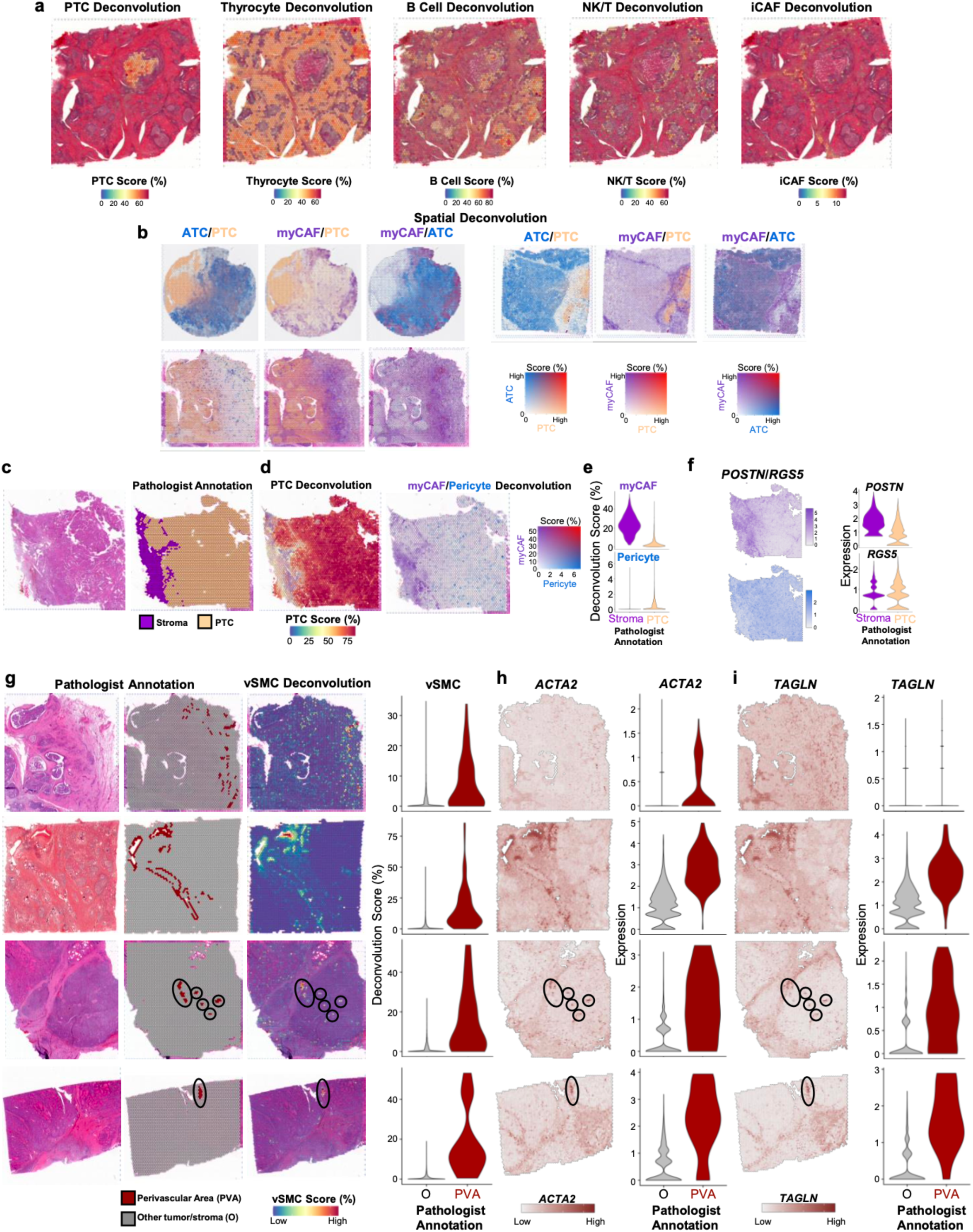
Spatial deconvolution of stromal populations. a, Spatial feature plots showing Robust Cell Type Decomposition (RCTD) of a pediatric papillary thyroid cancer (PTC) sample with background Hashimoto thyroiditis (Peds05). From left to right: PTC RCTD, thyrocyte RCTD, B cell RCTD, Natural Killer/T cell (NKT) RCTD, and inflammatory cancer-associated fibroblast (iCAF) RCTD. **b**, Spatial feature plots of three mixed PTC/anaplastic (ATC) samples (Thy1 Thy5, Thy10) showing ATC (blue), PTC (light orange), and myCAF (purple) RCTD scores. Mixing of deconvoluted populations is shown by a color gradient that becomes red. **c,** Hematoxylin and eosin staining of representative papillary thyroid cancer (PTC) Thy17 (left) with pathologist annotation of PTC (light orange) and stromal (purple) spatial barcodes (right). **d**, Spatial feature plots of representative PTC Thy17 showing Robust Cell Deconvolution (RCTD) of PTC (left, red color gradient) and myofibroblast cancer-associated fibroblast (myCAF)/pericyte (right, purple/blue color gradients) populations. **e**, Violin plots showing myCAF RCTD and pericyte RCTD scores within pathologist annotated stromal (purple) and PTC (light orange) barcodes. **f**, Spatial feature plots (left) and violin plots (right) depicting myCAF (*POSTN*) and pericyte (*RGS5*) marker gene expression in pathologist annotated stromal and PTC regions. **g**, Pathologist spatial annotation of large perivascular areas (PVAs) and spatial feature plot of vascular smooth muscle cell (vSMC) RCTD scores across four tumors. Left: hematoxylin and eosin staining and pathologist annotations. Middle: vSMC RCTD score plots. Right: violin plots of vSMC RCTD score stratified by barcodes whether spatial barcodes are in large PVAs. **h-i**, Spatial feature plots (left) and associated violin plots (right) for vSMC marker genes (**h**) *ACTA2* and (**i**) *TAGLN*. From top to bottom tumors are Thy5, Peds02, Thy9, Thy13 for **g-i**.

**Extended Data Fig. 7:**
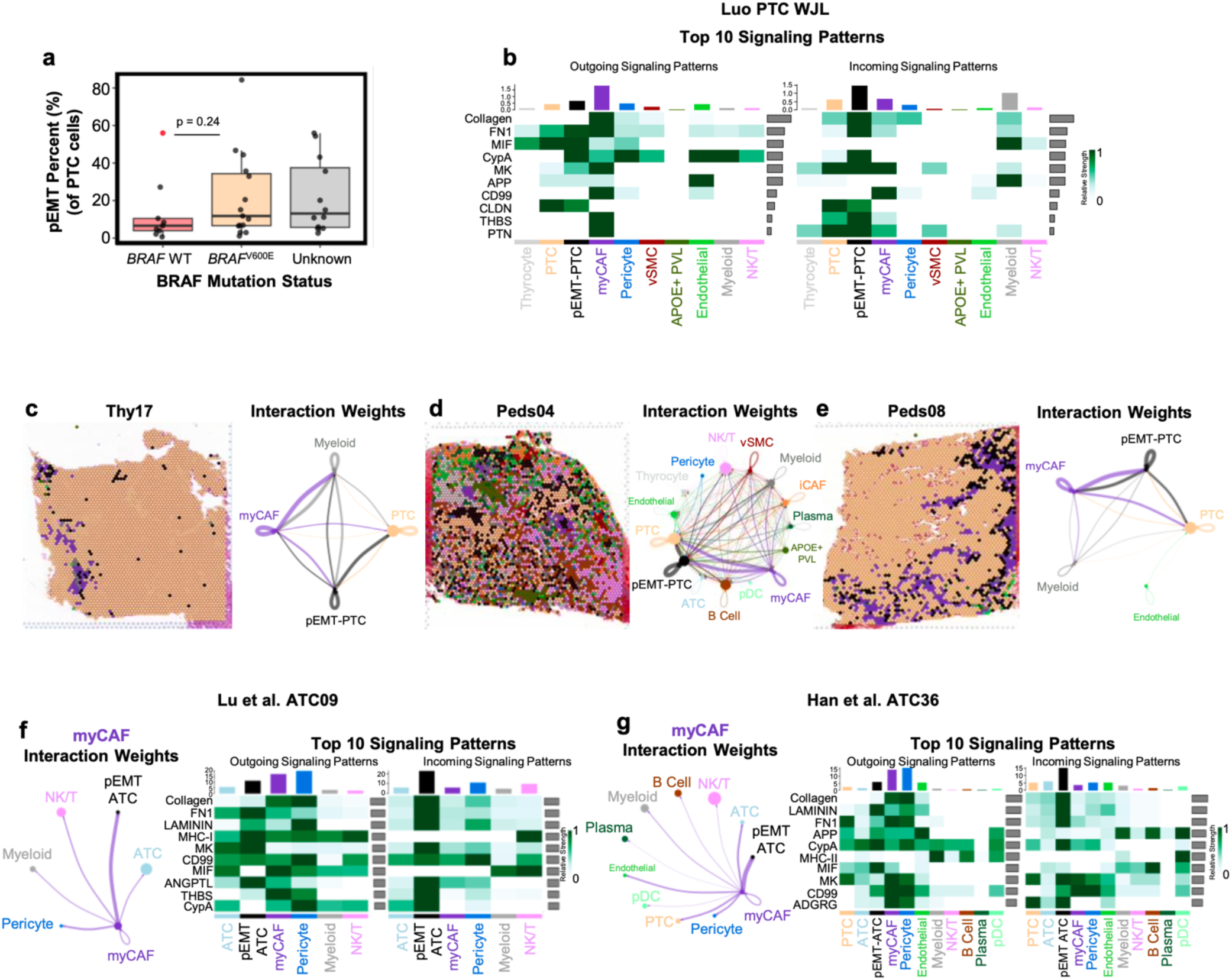
Ligand-receptor interaction analysis between tumor and stromal populations. a, Boxplot depicting the percent of partial epithelial-mesenchymal-transition (pEMT) cells in the broad papillary thyroid cancer (PTC) cluster for each single-cell sample with at least 100 PTC cells split by *BRAF* wild-type (WT, light red), *BRAF*^V600E^ (light orange), or unknown *BRAF* status. P-value between *BRAF* WT and *BRAF*^V600E^ calculated with Wilcoxon rank sum test. The red colored dot indicates a paratumor sample. **b,** Heatmap showing the top 10 signaling patterns in representative PTC WJL from Luo et al. with outgoing signal scores on the left and incoming signal scores on the right. Single-cell populations are labeled on the bottom. **c-e,** Spatial ligand-receptor interaction analysis at the invasive edge of PTC samples for (**c**) Thy17, (**d**) Peds04, and (**e**) Peds08. Left: Spatial plot with labeling of each spatial barcodes by the population with the highest Robust Cell Type Decomposition (RCTD) score in each sample. Right: spatial ligand-receptor interaction weights between labeled populations on the left. The width of lines indicates interaction weights. **f,g,** Ligand-receptor interaction analysis in *BRAF*^V600E^ anaplastic thyroid cancer (ATC) single-cell samples (**f**) Lu et al. ATC09 and (**g**) Han et al. ATC36 that have pEMT-ATC populations. Left: ligand-receptor interaction weights between single-cell populations. The width of lines indicates interaction weights. Right: The top 10 signaling patterns with outgoing signal scores on the left and incoming signal scores on the right. Single-cell populations are labeled on the bottom. Abbreviations: myCAF, myofibroblast cancer-associated fibroblast; NK/T; vSMC, vascular smooth muscle cell; PVL, perivascular-like; natural killer/T cell.

**Extended Data Fig. 8:**
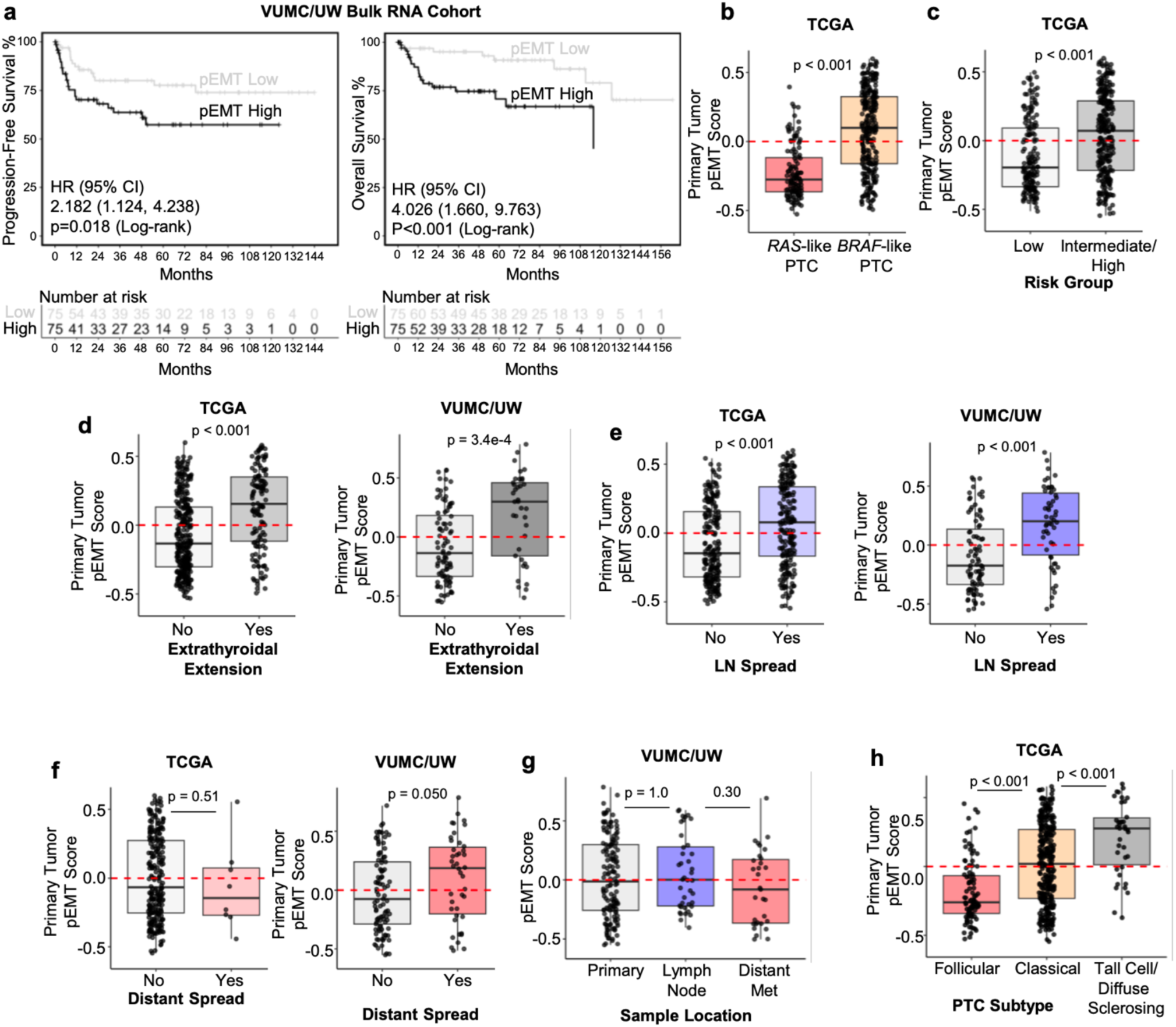
pEMT gene signature is associated with aggressive thyroid cancer tumor phenotypes. a, Progression-free survival (left) and overall survival (right) Kaplan-Meier curves for the Vanderbilt University Medical Center/University of Washington thyroid cancer bulk RNA-sequencing cohort (VUMC/UW bulk RNA cohort) for single-sample gene set variation analysis (ssGSVA) of a partial epithelial-mesenchymal transition (pEMT) gene list from Puram et al. split into pEMT-high and pEMT-low by 50^th^ percentile pEMT score. P-values calculated with log-rank test. **b-c,** Boxplots depicting primary tumor pEMT ssGSVA score in (**b**) The Cancer Genome Atlas (TCGA) papillary thyroid carcinomas (PTCs) stratified by *RAS*-like versus *BRAF*-like gene expression, (**c**) TCGA PTCs stratified by risk group as defined by the 2009 American Thyroid Association guidelines, (**d**) TCGA PTCs (left) and VUMC/UW malignant cohort (right) stratified by the presence or absence of extrathyroidal extension, (**e**) TCGA PTCs (left) and VUMC/UW malignant cohort (right) stratified by the presence or absence of lymph node spread, and (**f**), TCGA PTCs (left) and VUMC/UW malignant cohort (right) stratified by the presence or absence of distant metastasis. P-values for **b-f** calculated with Wilcoxon rank sum test. **g,** pEMT score in the VUMC/UW malignant cohort stratified by sample location. **h**, Primary tumor pEMT score for TCGA PTCs stratified by histology subtype. Tall cell and diffuse sclerosing PTC subtypes are grouped. P-values for **g-h** calculated with Wilcoxon rank sum test with Bonferroni correction.

**Supplementary Table 1:**
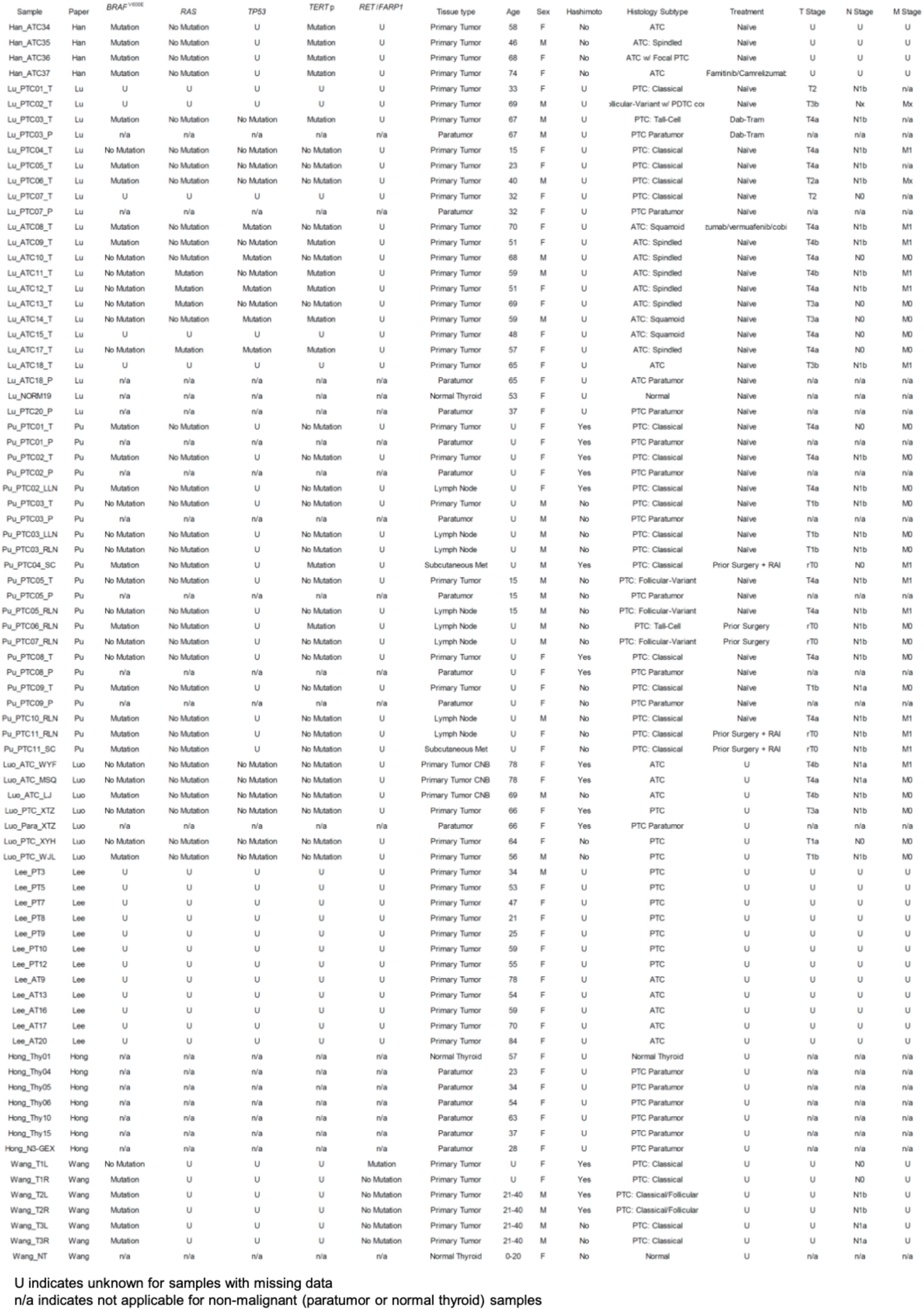
Single-cell atlas individual sample meta data.

**Supplementary Table 2:**
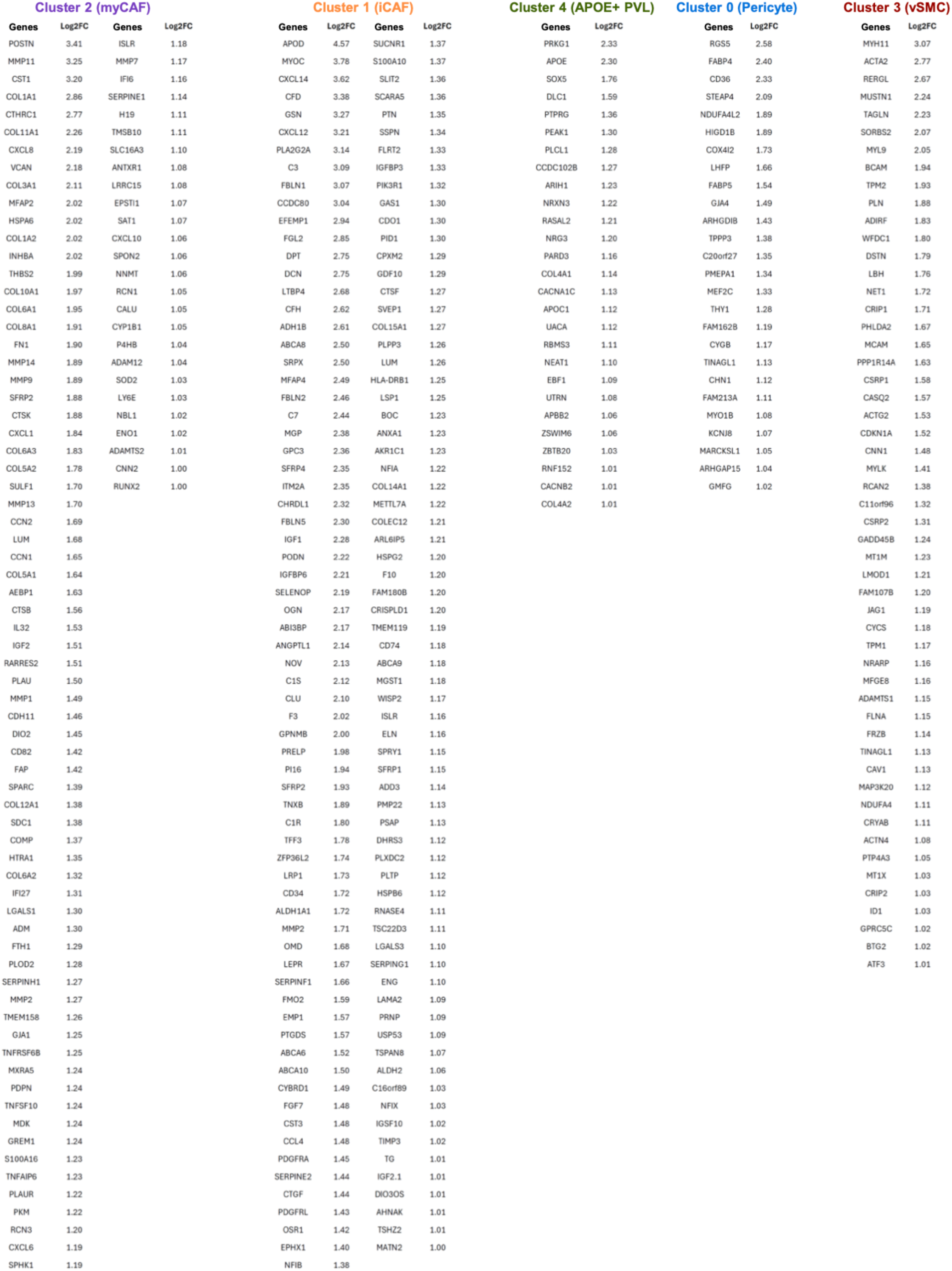
Marker genes of CAF and PVL populations.

